# HTLV-1 intasome recruits the PP2A-B56 holoenzyme and restricts its phosphatase activity

**DOI:** 10.64898/2026.05.01.722153

**Authors:** Jordan J Minnell, Emma Punch, Teresa Vanzo, Roghaiyeh Safari, Nora B. Cronin, Peter Cherepanov, Goedele N. Maertens

## Abstract

Integrase catalyses the insertion of a DNA copy of the retroviral genome into host cell chromatin. Human T-cell lymphotropic virus type 1 (HTLV-1) and other deltaretroviral integrases associate with protein phosphatase 2A holoenzymes containing B56 regulatory subunits (PP2A-B56). Here, we show that integrase mutants defective in binding to most B56 isoforms retain intrinsic DNA strand transfer activity but are impaired in establishing infection. Using single-particle cryo-EM, we determined the structure of the simian T-cell lymphotropic virus type 1 (STLV-1) intasome in a 0.5-MDa complex with two copies of the heterotrimeric PP2A-B56γ holoenzyme at 2.8 Å resolution. The structure reveals that, in addition to engaging B56, integrase forms direct contacts with the catalytic subunit of PP2A and sterically occludes the phosphatase active site, preventing substrate access. Consistent with these findings, we show that the HTLV-1 intasome suppresses PP2A catalytic activity in a manner dependent on the integrase LxxIxE short linear motif. We further demonstrate that pharmacological inhibition of PP2A does not impair HTLV-1 infection. Together, our results indicate that recruitment of PP2A-B56 by the intasome serves a structural rather than catalytic function.

## Introduction

Human T-cell lymphotropic virus type 1 (HTLV-1) is a deltaretrovirus that causes debilitating and often fatal disease. It is widespread in endemic regions, including parts of Japan, the Caribbean, sub-Saharan Africa, and South America [1], and there are currently no curative therapies available. A hallmark of the retroviral lifecycle is the integration of a DNA copy of the viral RNA genome into host chromatin. This reaction is catalysed by the viral integrase (IN) enzyme. Following reverse transcription, IN assembles on the viral DNA (vDNA) ends forming a synaptic complex referred to as the intasome or preintegration complex (PIC). IN then catalyzes (*i*) the 3’ processing reaction where a dinucleotide is removed from the 3’ long terminal repeat (LTR) ends producing nucleophilic 3’ OH ends, followed by (*ii*) the strand transfer reaction during which these nucleophiles attack the phosphodiester backbone of the target DNA across the major groove [2]. In case of HTLV-1, strand transfer takes place with a 6-bp stagger, which are repaired by host cellular enzymes, resulting in 6-bp duplication of target DNA sequence abutting the integration site. Whilst each retroviral species displays a defined consensus motif at the site of integration [3, 4], the integration site distribution is governed by multiple factors. This is best described for human immunodeficiency virus type 1 (HIV-1), a lentivirus, which depends on two host proteins, LEDGF/p75 and CPSF6, that interact respectively with IN and the viral capsid to integrate the proviral DNA into actively transcribed chromatin [5, 6]. Murine leukaemia virus (MLV) tethers its preintegration complex (PIC) via the p12^Gag^ protein to chromatin during cell division [7], after which the BET proteins take over and direct the PIC to transcription start sites, enriched with acetylation marks [8-10]. How HTLV-1 integration is modulated is currently unknown.

We have shown that HTLV-1, simian T-cell lymphotropic virus type 1 (STLV-1), and other deltaretroviral INs interact with PP2A-B56, a ubiquitous cellular Ser/Thr phosphatase that is a strong candidate for the role of the deltaretroviral PIC targeting factor [11]. PP2A comprises three subunits, the catalytic (C) and the scaffold (A), of which there are two highly conserved isoforms α and β, form the core enzyme. Binding of the PP2A core to any of the four families of structurally and functionally divergent regulatory subunits, B, B’/B56, B’’ or B’’’/STRIATIN forms the four major flavours of the PP2A holoenzymes. Substrate specificity and intracellular localization of PP2As are determined by their regulatory subunits. We have shown that B56 stimulates concerted integration activity of HTLV-1 (and -2) IN [11] and characterized the structure of the STLV-1 intasome in complex with B56 [12]; a homologous structure of the HTLV-1 strand transfer complex intasome was obtained by Aihara and colleagues [13]. We and others have shown that deltaretroviruses employ molecular mimicry to engage with PP2A-B56 [12-14]. Accordingly, deltaretroviral INs bind B56 in a manner similar, but not identical, to numerous endogenous PP2A–B56 interactors that contain the LxxIxE short linear motif (SLiM) [15, 16]. SLiMs are typically found within intrinsically disordered regions, where they mediate protein-protein interactions and are often subject to post-translational modifications [17]. Deltaretroviruses are not unique in targeting PP2A, and all three subunits of the heterotrimeric phosphatase have been shown to be hijacked by viral factors [18].

Using a combination of fluorescence polarization, pull-down and immunoprecipitation we characterized mutants of HTLV-1 IN and B56. We identified catalytically active IN SLiM mutants that are defective for binding to B56. HTLV-1 viruses carrying these mutations are highly defective for infectivity. Finally, using cryo-EM, we refined the structure of the intasome in complex with the PP2A-B56 holoenzyme and showed that the intasome blocks the PP2A active site. Using phosphatase assays we confirm that IN, in a SLiM-dependent manner, inhibits PP2A-B56 activity, suggesting that the deltaretroviral DNA integration machinery recruits PP2A-B56 for structural reasons rather than for its phosphatase activity.

## Results

### The B56-specific LxxIxE SLiM is critical for the HTLV-1 IN:B56 interface *in vitro*

Recently, we showed that alanine substitutions of the canonical STLV-1 IN SLiM residues (L213, I216 and E218) resulted in a loss of B56γ(11-380) binding [12]. Using fluorescence polarization (FP), we confirmed that the structurally equivalent residues of HTLV-1 IN are critical for the interaction with B56γ (Supplementary Table S1). Reciprocal mutations of B56γ residues involved in SLiM recognition (H187A and R197A) completely abrogated binding to peptides of both wild type (WT) HTLV-1 IN and RepoMan, an endogenous PP2A-B56 substrate [15]. Moreover, alanine substitutions of the B56γ residues lining up its isoleucine binding pocket (I227, I231, H187 and Y190) [16] significantly weakened the interactions. By contrast, and in agreement with our previous results [11], alanine substitution of B56γ R188 – a residue important for binding phosphorylated endogenous substrates by PP2A-B56 [16] – did not significantly affect the interaction with either peptide (Supplementary Figure S1b,c).

Next, we used hexahistidine (His_6_) pull-down assays to assess the requirement of the B56γ SLiM-binding site for the interaction with full-length HTLV-1 IN. Consistent with the FP assay results, His_6_-tagged HTLV-1 IN efficiently pulled down WT B56γ(11– 380), but not its H187A or R197A variants (Figure 1a). The interaction was reduced by alanine substitutions at residues I227, I231, and Y190, whereas the R188A substitution had no detectable effect (Figure 1a). The reciprocal experiment was conducted using alanine point mutants of the HTLV-1 IN SLiM residues: L213A, I216A and E218A. As expected, each mutant reduced binding to B56γ(11-380): L213A and I216A resulted in 27 and 31-fold reductions in binding compared to the WT, the double mutant L213A/I216A displayed a 10-fold reduction in binding compared to WT, whilst E218A completely abolished the interaction (Figure 1b).

**Figure 1.**
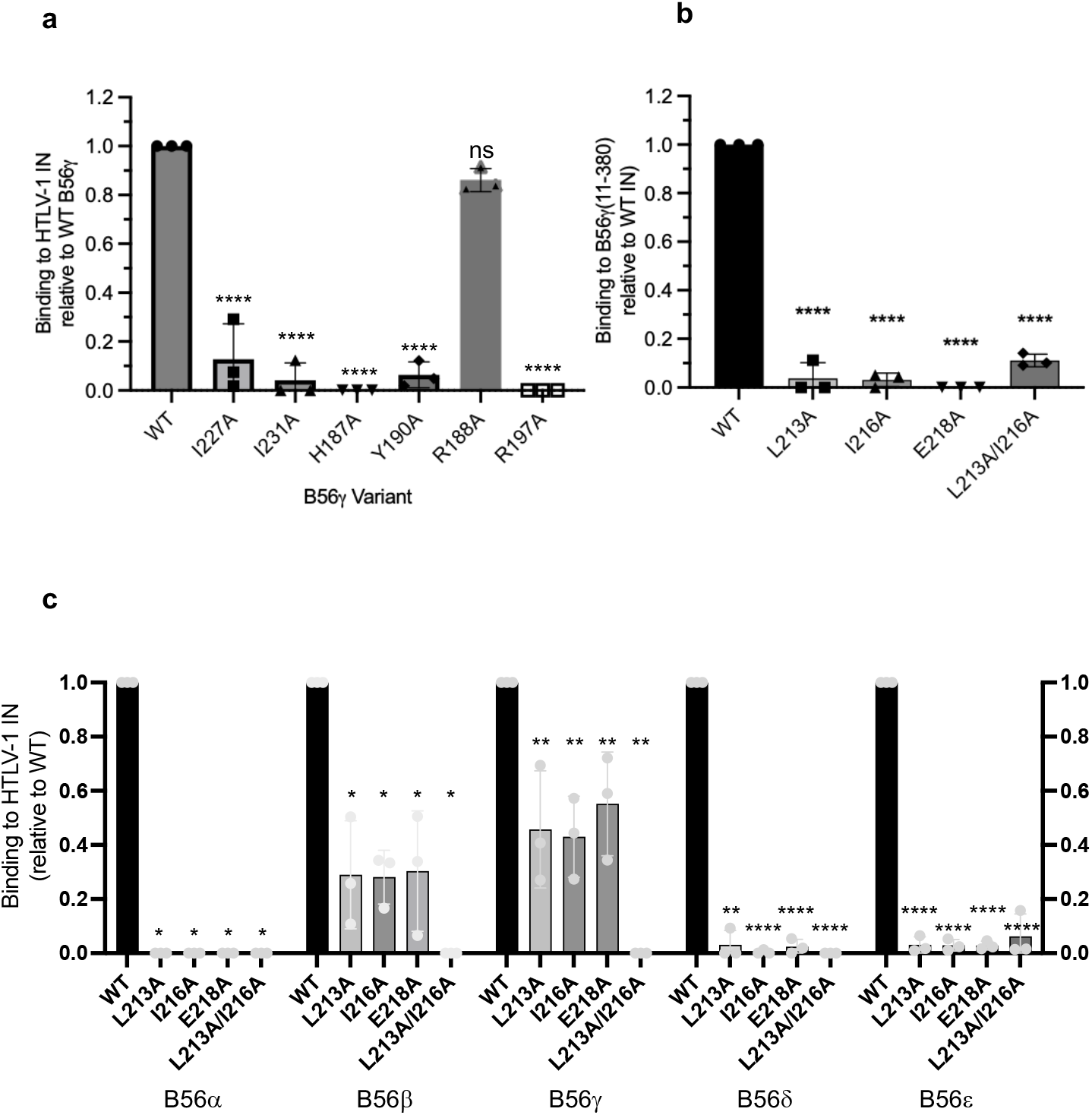
Binding of B56 isoforms to HTLV-1 IN SLiM mutants. ***a***, His_6_-tag pull-down using wild type His_6_-tagged HTLV-1 IN as bait and B56γ(11-380) WT and single point mutants as prey. Relative binding compared to WT B56γ(11-380) is shown. ***b***, His_6_-tag pull-down using WT B56γ(11-380) as prey and WT or mutant His_6_-tagged HTLV-1 IN as prey as indicated on the graph. Relative binding compared to WT HTLV-1 IN is shown. ***c***, Flag-immunoprecipitation of Flag-tagged WT or mutant HTLV-1 IN in 293T cells. Endogenous B56α-δ and HA-tagged B56ε were detected by western blot. Relative binding compared to WT HTLV-1 IN was quantified by densitometry. Averages and standard devations of three independent immunoprecipitations or pull-downs are shown.

The previous experiments were conducted *in vitro* using recombinant full-length HTLV-1 IN and a B56γ fragment that is sufficient for WT binding. Human cells express five major B56 isoforms (α-ε), each further diversified through alternative splicing and alternative translation [19, 20]. Therefore, we wanted to investigate how the introduction of the SLiM mutations in HTLV-1 IN affect binding to all five endogenous B56 family members. Flag-tagged full-length HTLV-1 IN was transiently overexpressed in the human embryonic kidney 293T cells, and was immunoprecipitated (IP) from the cell lysate using an anti-Flag antibody. Western blotting of the IP fraction revealed the IN mutants produced a significant but varied reduction in recovery of all B56 family members tested (Figure 1c, Supplementary Figure S2). The other PP2A regulatory family members, B55, PR72 and STRIATIN, were undetectable in the IP fraction, corroborating our previous observations that the interaction is specific to the B56 family members [11]. Each of the four tested IN SLiM mutations abolished the binding to B56α and B56δ (Figure 1c), whilst only about 30% and 50% of B56β and B56γ, respectively was recovered with either of the Flag-IN SLiM mutants compared to WT Flag-IN (Figure 1c). We could not detect endogenous B56ε from 293T cell lysate by Western blotting. We therefore transiently co-expressed the Flag-tagged IN proteins together with haemagglutinin (HA)-tagged B56ε, which afforded the detection of the interaction. By contrast, neither of the IN SLiM mutants are able to bind HA-B56ε (Figure 1c). Collectively, these results demonstrate that HTLV-1 IN interacts with all human B56 isoforms, and that this interaction depends on the SLiM and the corresponding SLiM-binding region in B56.

### HTLV-1 IN SLiM mutants are catalytically active

Before evaluating functionality of HTLV-1 IN SLiM mutants in the context of virus infectivity, we wanted to ensure that the amino acid substitutions did not abrogate catalytic activity of the enzyme. To this end, we produced full-length, untagged HTLV-1 IN variants and tested their *in vitro* strand-transfer activities. This assay, described previously [11, 21-23], uses short double-stranded oligonucleotides that mimic the ends of viral DNA (vDNA) and a supercoiled plasmid as the target DNA. Concerted strand transfer of a pair of short donor DNA molecules results in linearization of plasmid DNA, whereas half-site processes (integration of a single donor molecule into one DNA strand of the target plasmid) results in accumulation of relaxed DNA circles. Both types of DNA products can be detected and quantified following separation in agarose gels. We observed no inhibitory effect of the SLiM mutations on the intrinsic catalytic activity of HTLV-1 IN; in fact, all four mutants exhibited moderately enhanced strand transfer activity compared to WT, with the L213A/I216A double mutant showing ~1.7-fold higher activity (Fig. 2a). By contrast, the catalytic-site mutant D122N displayed no detectable strand transfer activity.

**Figure 2.**
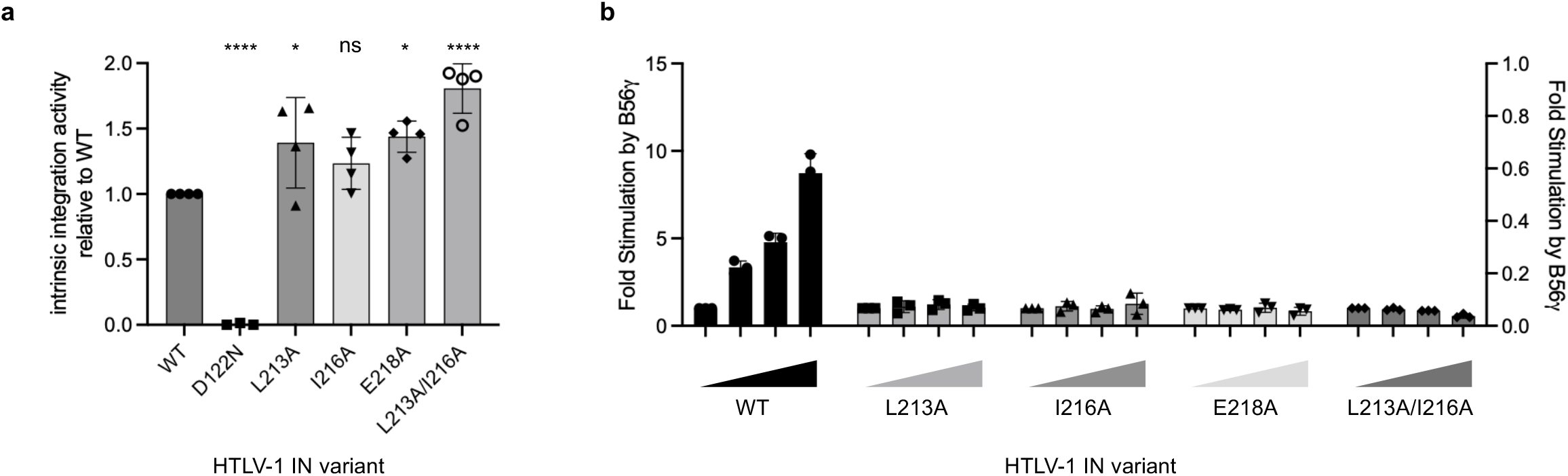
HTLV-1 IN SLiM mutants are catalytically active. ***a***, Relative intrinsic HTLV-1 IN activity of SLiM mutants compared to WT HTLV-1 IN. The catalytically dead (IN D122N) was included as negative control. Average and standard deviations of four independent assays are shown. ***b***, HTLV-1 IN activity is stimulated by addition of WT B56γ(11-380), HTLV-1 IN SLiM mutants unable to bind to B56γ(11-380) are not stimulated upon addition of increasing amounts of B56γ(11-380). Average and standard devaiations of three independent assays are shown.

We have shown that B56γ stimulates deltaretroviral IN strand transfer activity *in vitro* [11]. Concordantly, addition of B56γ(11-380) to WT HTLV-1 IN stimulated strand transfer in a dose-dependent manner (Figure 2b). As expected, the catalytic activity of each IN SLiM mutant tested was not enhanced by B56γ(11-380) (Figure 2b). These results provide point mutants that both abrogate B56 binding but have no adverse effect on the intrinsic enzymatic function of IN.

### The IN SLiM is essential for HTLV-1 infectivity

To investigate the significance of the IN : B56 interaction during HTLV-1 infection, we introduced the IN SLiM point mutations in the integration-dependent reporter system developed by Derse and colleagues [24]. The catalytic site mutant D122N was included as a control to estimate background GFP signal arising from unintegrated DNA and inadvertent recombination events. The WT HTLV-1 vector was able to convert ~1.5% of target cells to GFP-positive state. Such relatively low levels of infectivity are typical for tissue culture experiments using HTLV-1 proviral constructs [13, 24]. In this single round reporter assay, all SLiM mutants tested were unable to establish infection, since either no GFP^+^ cells could be detected, or the signal was similar to HTLV-1 IN D122N mutant (Figure 3a,b). To further characterise the impact of the IN SLiM mutations on HTLV-1 infection, we generated replication-competent viral particles by transfecting HEK293T cells with the pACH molecular clone [25] and assessed their infectivity in a co-culture assay. We produced WT HTLV-1, HTLV-1 IN^D122N^ (IN active site mutant defective for integration), along with a panel of HTLV-1 IN SLiM mutants, as well as pACH^ΔENV^. The latter construct, carring a deletion in the viral *env* gene, served as an additional negative control, as functional viral envelope glycoprotein is essential for HTLV-1 entry [26]. Western blotting of transfected cells using a range of antibodies revealed expression of HTLV-1 Tat, Gag and gp46 proteins (Figure 3c). Moreover, Gag was detected in the concentrated viral supernatant from each producer cell line using HTLV-1 capsid-specific antibodies (Figure 3c). Viral reverse transcriptase (RT) present in the supernatants was quantified by SYBR green I-based product-enhanced RT (SG-PERT) assay [27], and the measured RT levels used to normalise the viral inputs (Supplementary Figure S3a). The infection experiments were done by co-culture of the transfected HTLV-1 producing 293T cells with a human osteosarcoma (HOS) cell line harbouring a Tax-inducible GFP reporter construct [28]. Upon infection of the indicator cells, Tax expressed from the integrated provirus activates the minimal HTLV-1 LTR promoter resulting in GFP expression. Intriguingly, we noted transactivation of GFP expression in the target cells during infection with the D122N IN-defective virus as early as 72 h post-infection, as measured by flow cytometry (Supplementary Figure S3b). This was not observed from infection with HTLV-1 ΔEnv control although both pACH mutants were capable of expressing Tax (Figure 3c), suggesting that early post-infection, Tax is likely expressed from unintegrated viral DNA. The percentage of GFP^+^ HOS cells was analysed by flow cytometry 9-10 days post-infection to allow sufficient time for the decay of non-integrated viral DNA. All HTLV-1 IN mutant viruses were significantly less infectious compared to WT virus; both the HTLV-1 IN^L213A^ and IN^E218A^ mutant viruses displayed ~4-fold reduction in infectivity, whilst the HTLV-1 IN^I216A^ mutant was as defective as the IN active site mutant D122N (Figure 3d). These results demonstrate that an intact IN SLiM is critical for HTLV-1 infectivity.

**Figure 3.**
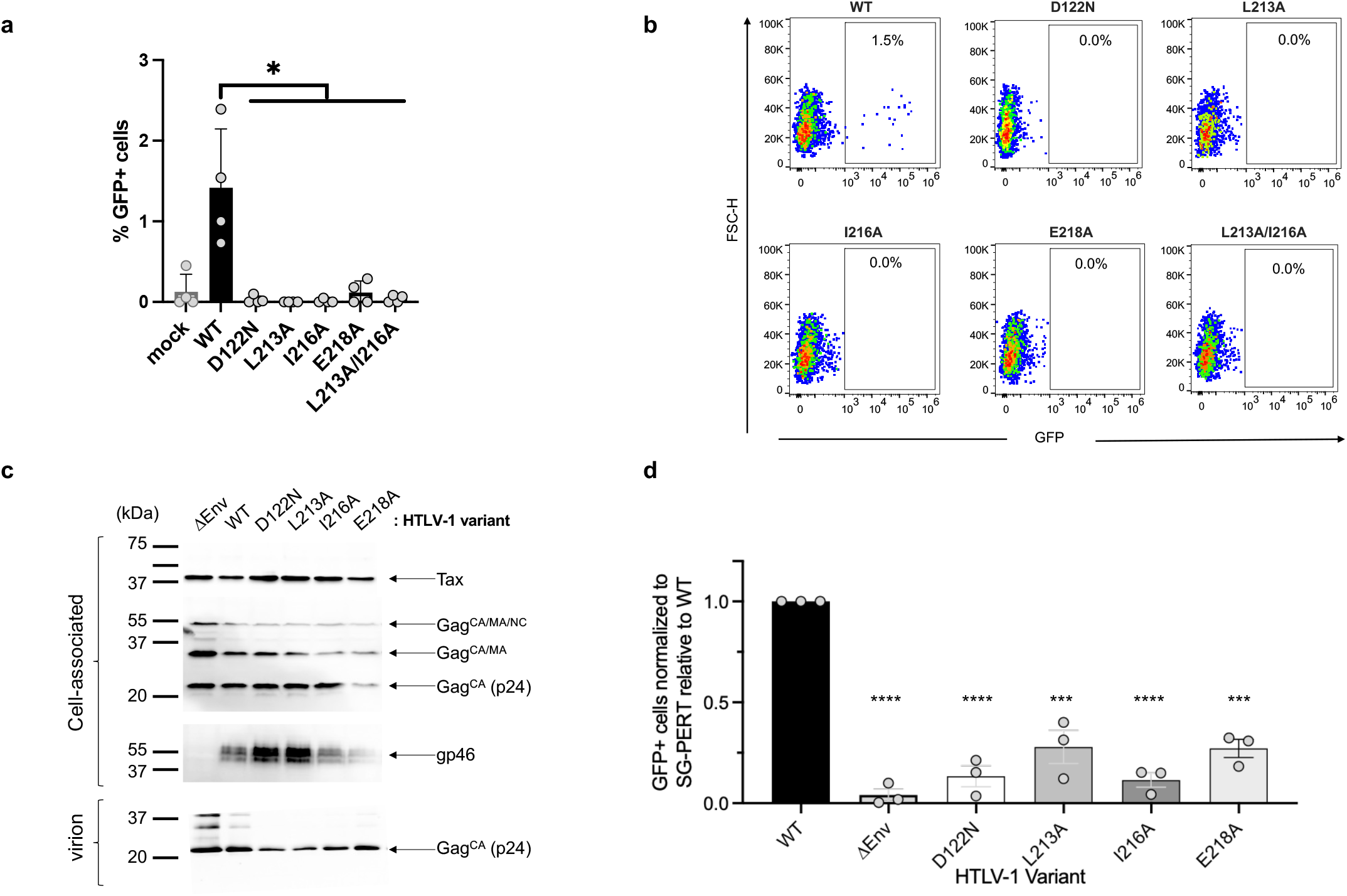
Mutation of the HTLV-1 IN SLiM abrogates HTLV-1 infectivity. ***a***, Bar chart showing infection efficiency of single-round HTLV-1 GFP-reporter vector, WT, catalytically dead (D122N), and SLiM mutants. Averages and standard deviations of four independent replicates are shown. ***b***, Representative example of FACS data illustrating infection efficiency (% GFP positive cells) of WT, catalytically dead (D122N) and four HTLV-1 IN SLiM mutant following single round infection using a GFP-reporter vector. ***c***, Western blots illustrating expression of cell-associated HTLV-1 Tax, p24^Gag^, gp46, and virion-associated p24. Antibodies used are indicated on the right, molecular weight markers on the left. HTLV-1 variants are indicated at the top of the blots. ***d***, Relative infection efficiency normalized to WT HTLV-1 using WT, catalytically dead (D122N) and IN SLiM mutants. An additional negative control in which the ENV gp46 was removed (DEnv) was included. Infection was measured by FACS, measuring GFP fluorescence in the HOS-GFP reporter cell line following 10 days of infection. Averages and standard deviations of three independent biological repeats are shown.

### Cryo-EM structure of the STLV-1 intasome in complex with PP2A-B56 holoenzyme

B56γ residues 11-380 are minimally sufficient for interaction with HTLV-1 and STLV-1 INs, and we previously characterised the STLV-1 intasome in complex with this B56 fragment [12]. However, in cells, deltaretroviral INs engage with the full ~158 kDa heterotrimeric PP2A holoenzyme [11, 12]. To assess whether the STLV-1 intasome can accommodate the complete phosphatase complex, we analysed IN-DNA nucleoprotein assemblies formed in the presence or absence of the PP2A catalytic and scaffold subunits using electrophoretic mobility shift assays (EMSA). Using fluorescently labelled oligos mimicking vDNA ends, we assembled the STLV-1 intasome in complex with full-length B56γ, as described previously for the minimal B56γ fragment, and subsequently supplemented the complexes with full-length human PP2A scaffold (Aα) and catalytic (Cα) subunits. Electrophoretic separation under non-denaturing conditions revealed a reproducible shift of the intasome band in the presence of the PP2A core components (Supplementary Figure S4a), indicating complex formation. The assembled complexes remained stable for at least 2 h on ice (Supplementary Figure S4a). For structural analysis, the intasome-B56γ complex was purified by size-exclusion chromatography and subsequently supplemented with PP2A scaffold and catalytic subunits. Negative-stain EM revealed distinct particles with a maximal dimension of ~28 nm, approximately twice the size of the minimal STLV-1 intasome described previously [12] (Supplementary Figure Figure S4b,c). The characteristic horse-shoe shaped density of the PP2A Aα scaffold was evident at both ends of the extended intasome envelope (Supplementary Figure S4b).

To characterize the structure of the intasome - PP2A-B56γ holoenzyme complex, we vitrified the complex on graphene oxide support and imaged by cryo-EM (Supplementary Figure S5a). Single-particle analysis yielded a three-dimensional reconstruction of the twofold symmetric assembly at an overall resolution of 2.8 Å (Supplementary Fig. S5b, Supplementary Table S2). The intasome, along with two tightly associated copies of the complete PP2A holenzyme (Figure 4a,b) were well defined in the cryo-EM map (Supplementary Figure S6). Local resolution ranged from 2.5 to 3.2 Å across most of the assembly, including the four IN subunits, the pair of vDNA ends, and all three PP2A subunits (Supplementary Figure S5b). To facilitate model building, we performed a local reconstruction of one half of the complex, which improved the overall resolution to 2.6 Å and enhanced the definition of the peripheral regions of the scaffold subunit (Supplementary Figure S5c). Both copies of the PP2A heterotrimer within the complex are structured very similar to the holoenzyme in isolation (Supplementary Figure S7), suggesting that the intasome can recruit the preassembled heterotrimeric phosphatase.

**Figure 4.**
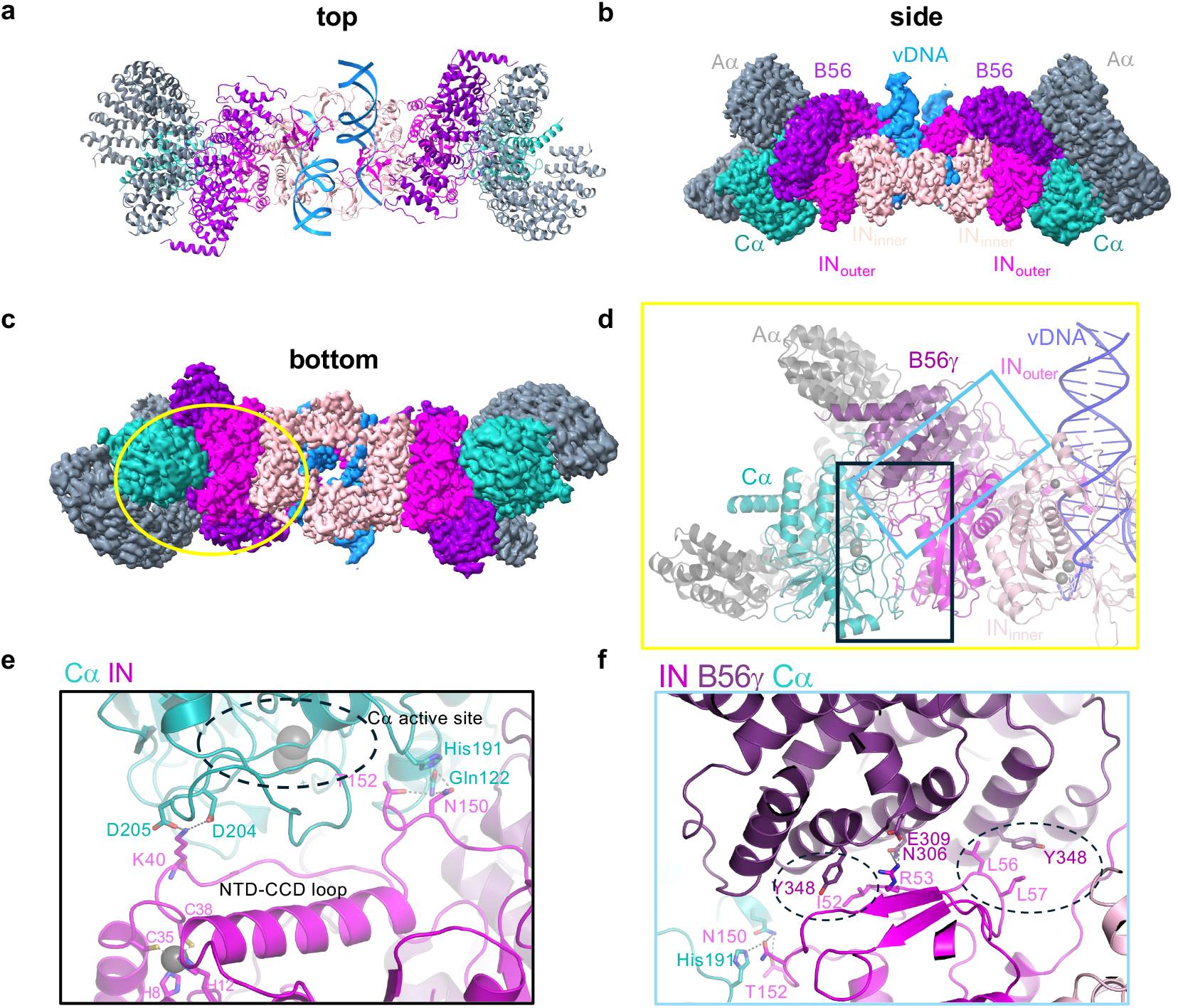
Cryo-EM structure of STLV-1 intasome : PP2A-B56γ holoenzyme. ***a***, The intasome : PP2A-B56γ holoenzyme structure. The synaptic complex comprises a tetramer of IN (two inner IN (light pink) and two outer IN (magenta) molecules) that synapse two viral DNA (vDNA, blue) ends together. On either side of the intasome, the PP2A-B56γ holoenzyme is bound. The scaffold (Aα) is shown in grey, catalytic subunit (Cα) in cyan and B56γ in purple. ***b***, Side view of the 2.8 Å cryo-EM map reconstruction overlaid on top of the model. ***c***, bottom view of the cryo-EM map. Yellow circle indicates area of interest magnified in panel d. ***d***, zoom into the expanded interaction interface between the outer IN and B56γ, and IN and Cα. Individual chains are indicated. ***e***, interaction interface between IN and Cα. Hydrogen bonds and salt bridges are shown with dashed lines. The Cα active site (with two bound Mg^2+^ ions shown as grey spheres) is indicated with a dashed circle. Residues chelating Zn^2+^ (grey sphere) in the IN/NTD are shown as sticks. ***f***, Interaction interface between IN and B56γ, hydrogen bonds and salt bridges are shown with dashed lines. Hydrophobic interactions are highlighted in the dashed circles. Panels ***a-c*** were made using ChimeraX, panels ***d-f*** were made with Pymol 3.1.6.1.

The bulk of the intasome-PP2A contact surface is formed in the IN-B56γ interface, measuring ~2,200 Å^2^, of which ~ 1,900 Å^2^ is contributed by the outer IN subunit (chain A). This interface closely resembles that observed in the 3.3-Å cryo-EM structure of the intasome bound to the minimal B56γ fragment [12], but is resolved here at substantially higher resolution (Supplementary Figure S5). The interaction is centred on the conserved SLiM-binding groove of B56γ, which engages the IN SLiM, and is further stabilised by additional contacts that extend beyond the canonical motif-binding pocket, reinforcing the IN-B56γ association.

The dominant IN–B56γ interaction orients the PP2A holoenzyme such that the catalytic site of the PP2A Cα subunit faces the outer IN chain. In contrast to the extensive IN-B56γ interface, the IN–Cα contact is relatively limited (~500 Å^2^). STLV-1 IN residues 39-53 form a linker connecting the N-terminal domain (NTD) and the catalytic core domain (CCD). As observed in other retroviral intasomes, the NTD–CCD linkers of the inner IN subunits (corresponding to chains B and G in the refined model) are well ordered and play integral roles in stabilising the intasome architecture [12, 13, 29-31]. By contrast, the corresponding linkers of the outer IN subunits (chains A and F) are positioned directly in front of the PP2A active sites (Fig. 4c, d). This disordered loop is stabilized on either side by IN residues Lys40, Asn150 and T152 that directly associate with side chains of Cα residues Asp204, Asp205, Gln122 and His191, respectively (Figure 4e); whilst IN residues Ile52, Arg53, and His60 further stabilize the interaction via hydrogen bonds with B56γ residues Asn306, Glu309 and Glu310. Hydrophobic interactions between IN residues Ile52 and B56γ Tyr348, and IN Leu56 and Leu57 with B56γ Tyr269 further anchor the IN/CCD in place (Figure 4f).

Notably, the entire STLV-1 IN NTD-CCD linker is devoid of serine or threonine residues that when phosphorylated could serve as PP2A substrates (Figures 4e and 5a), arguing against the intasome acting as a substrate for the phosphatase. Instead, the positioning of the linker resembles an active-site occluding structural element observed in the complex of PP2A-B55 with the inhibitory protein ARPP19 (Figure 7; see Discussion), suggesting that the intasome sterically blocks substrate access and thereby inhibits phosphatase activity.

**Figure 5.**
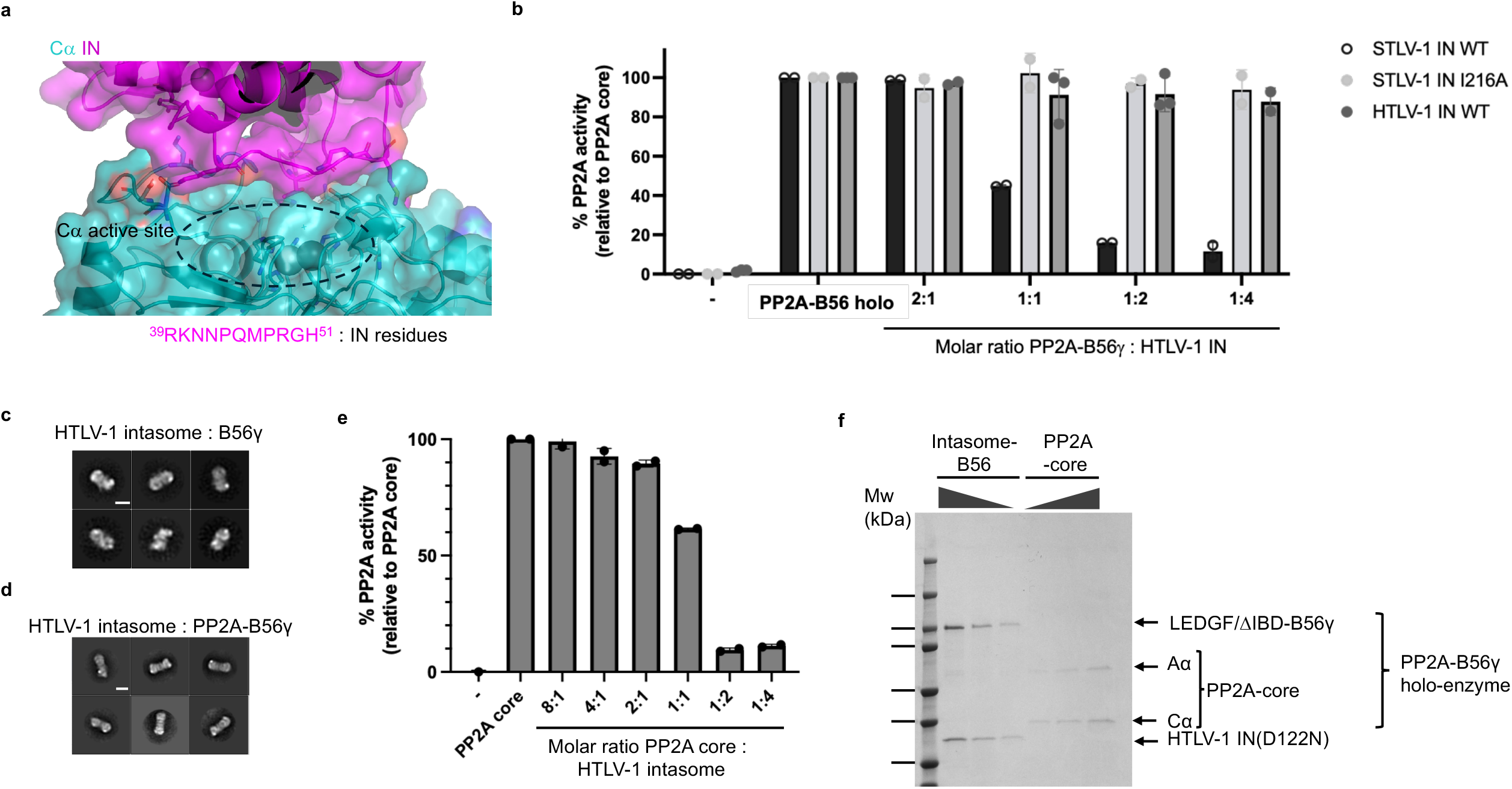
The HTLV-1 intasome blocks PP2A-B56 activity. ***a***, Surface view of the interaction interface between IN and Cα. Interacting residues on either side of the disordered loop are shown as sticks. Sequence of the disordered peptide is shown below the image. ***b***, Relative PP2A activity of PP2A-B56 alone, in the presence of increasing amounts of HTLV-1 IN, STLV-1 IN or the SLiM mutant STLV-1 IN(I216A). As a negative control (-) phosphatase inhibitors were added prior to adding the substrate. Averages and standard deviations of two independent biological replicates are shown. ***c***, 2D class averages of the HTLV-1 intasome : B56γ, and ***d***, HTLV-1 intasome : PP2A-B56γ. Scale bars in panel ***c*** and ***d*** are 10 nm. ***e***, Relative PP2A-B56 activity in the presence of increasing amounts of HTLV-1 intasome. Averages and standard deviations of two biological replicates are shown. ***f***, SDS-PAGE gel stained with colloidal Coomassie illustrating the complexes used in panel ***e*** loaded at different dilutions. Molecular weight markers are shown on the left, proteins are indicated on the right. Complexes loaded on gel are indicated above the gel.

### PP2A activity is blocked by the HTLV-1 intasome, and pharmacological inhibition of PP2A does not impact HTLV-1 infectivity

We previously reported that HTLV-1 IN did not inhibit PP2A holoenzymes co-purified with Flag-tagged B56γ from HeLa cells [11]. Here, we examined the catalytic activity of recombinantly purified PP2A-B56γ in the presence of STLV-1 IN, buffer alone, or a phosphatase inhibitor as negative control. By contrast to HTLV-1 IN, STLV-1 IN efficiently inhibited PP2A activity (Figure 5b) and this effect was dependent on the IN-B56γ interaction, as the IN SLiM mutant I216A failed to block phosphatase activity (Figure 5b). In agreement with our previous observations, pre-incubation with HTLV-1 IN reduced PP2A activity by only ~20%, even at fourfold molar excess (Figure 5b) [11]. STLV-1 IN binds B56γ with higher affinity than HTLV-1 IN [32], which may account for its greater inhibitory potency. Notably, these experiments were performed with free IN, rather than IN assembled within the intasome, where multivalent engagement of B56γ is expected to further stabilise the interaction and orient the catalytic subunit in the way observed in the cryo-EM structures. Indeed, as shown previously, free IN associates with B56γ in an equimolar ratio and is driven by the IN SLiM binding to the canonical SLiM binding pocket on B56γ [12]. In the context of the intasome, two IN molecules associate with one B56γ molecule; the outer IN SLiM associates with the canonical SLiM binding pocket, whilst the inner IN SLiM binds to the non-canonical SLiM binding site on B56γ [12].

Thus, we assembled the HTLV-1 intasome employing a similar approach as for the STLV-1 intasome, using recombinantly purified HTLV-1 IN D122N (catalytically inactive), LEDGF/ΔIBD-B56γ and DNA oligonucleotides mimicking the DNA product of strand transfer of the vDNA with target DNA [13]. HTLV-1 IN strand transfer (STC)-B56γ complexes were purified by size exclusion chromatography. Characterisation by negative stain-EM, in the presence and absence of the PP2A core enzyme, confirmed proper assembly of the holoenzyme complex (Figure 5c, d). Importantly, the HTLV-1 intasome efficiently inhibited PP2A-mediated dephosphorylation of a model phosphopeptide substrate in a dose-dependent manner (Figure 5e,f).

Given that IN blocks the active site of PP2A-B56, it is likely that the catalytic activity of the phosphatase is not required for infectivity. To investigate this, we pre-treated Jurkat cells with PP2A inhibitors (PP2Ai) okadaic acid (OA) or Rubratoxin-A (RubA), or DMSO (vehicle) and investigated the effect on HTLV-1 infection. As a negative control, we treated Jurkat cells with raltegravir (RAL), a potent IN strand transfer inhibitor that blocks HTLV-1 infectivity in tissue culture [23, 33]. Cells were treated with vehicle/PP2Ai prior to infection and during the course of co-culture with the persistently infected MT-2 cell line. Following depletion of MT-2 cells using magnetic beads, the PP2A inhibitors were removed and further integration were blocked by addition of RAL. High concentration of RubA (2 μM) and OA (20 nM) affected cell viability (Figure 6a), whilst the lower concentrations (10 nM OA, 600 nM RubA) did not. Importantly, neither of the PP2A inhibitors affected HTLV-1 infection, as measured by proviral DNA content (Figure 6b).

**Figure 6.**
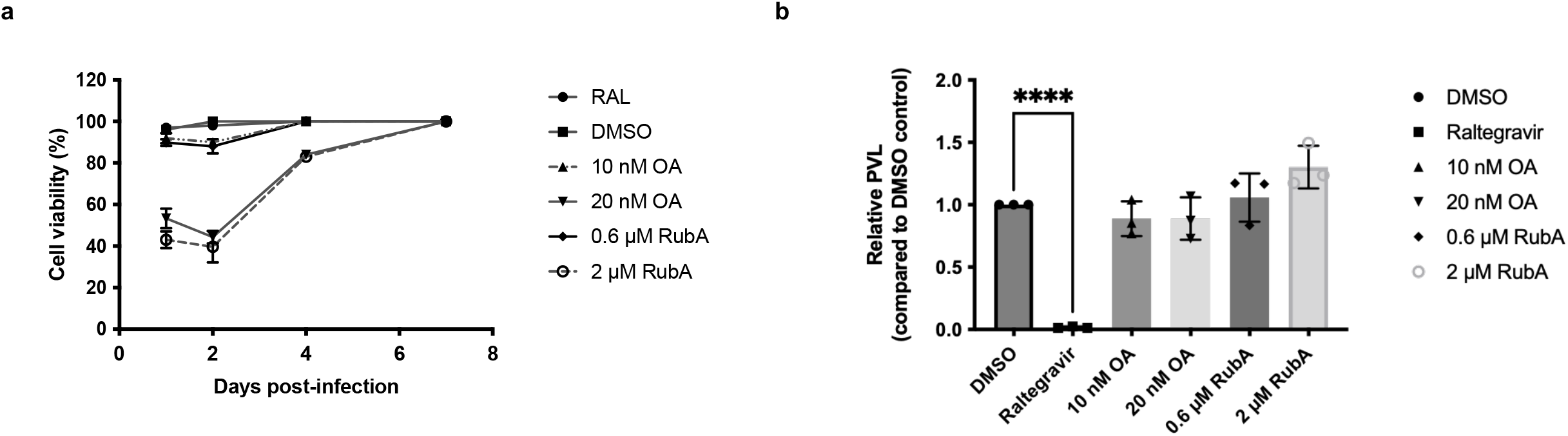
Inhibition of PP2A does not impact HTLV-1 infection efficiency. ***a***, Cell viability measured in Jurkat T cells following treatment with vehicle (DMSO), 10 mM raltegravir (RAL), 10 or 20 nM okadaic acid (OA) and 600 or 2000 nM rubratoxin A (RubA). ***b***, Relative HTLV-1 infection efficiency compared to DMSO control, measured by PVL. Averages and standard deviations of 3 independent biological replicates are shown.

## Discussion

Here, we have characterized the interaction between deltaretroviral INs (HTLV-1 and STLV-1) and the PP2A-B56 holoenzyme. We have found that interaction between the IN SLiM motif and B56 is critical to establish infection since mutation of the IN SLiM does not affect intrinsic IN catalytic activity yet abrogates B56 interaction and blocks HTLV-1 infection. Inhibiting PP2A using either okadaic acid or rubratoxin A did not reduce HTLV-1 infection under the conditions tested (Figure 6), supporting the structural model that PP2A-B56 is recruited primarily for a non-catalytic role. Indeed, we showed that in the context of the intasome HTLV-1 IN sterically blocks the Cα active site and inhibits its catalytic activity in a phosphopeptide-based assay. While, to our knowledge, this is the first case of a viral protein blocking the active site of PP2A-B56, various cellular factors are known to inhibit the phosophatase [34]. For example, cAMP regulated phosphoprotein 19 (ARPP19), an intrinsically disordered protein, inhibits PP2A-B55 activity during mitotic entry [35]. ARPP19 makes extensive contacts with both B55 and Cα. Superimposition of our structure with the PP2A-core enzyme bound to ARPP19 illustrates that STLV-1 IN residues 39-46 occupy a similar space as α4 of ARPP19 that blocks the Cα active site (Figure 7). The binding mode is not exactly the same, ARPP19 blocks the active site with a structured alpha-helix, whilst STLV-1 IN uses an intrinsically disordered loop held in place at either end. It is possible that the conformational flexibility of this intrinsically disordered peptide serves as a versatile platform for transient interactions with diverse other binding partners that have yet to be identified. Notably, phosphorylation of ARPP19 is not required to bind to PP2A-B55, but the phosphorylated form (pSer62) is a more potent inhibitor [35]. Residues in the extended NTD-CCD linker are less conserved in HTLV-2 and bovine leukaemia virus (BLV) IN. In contrast to the other deltaretroviral INs, HTLV-2 IN Ser43, and BLV IN Thr44 could be subject to phosphorylation and align well with Ser62 of ARPP19. It is possible that throughout evolution [36], primate T-lymphotropic viruses 1 and 3 (and their zoonotic progeny) have lost the need to depend on phosphorylation to interact with PP2A-B56.

**Figure 7.**
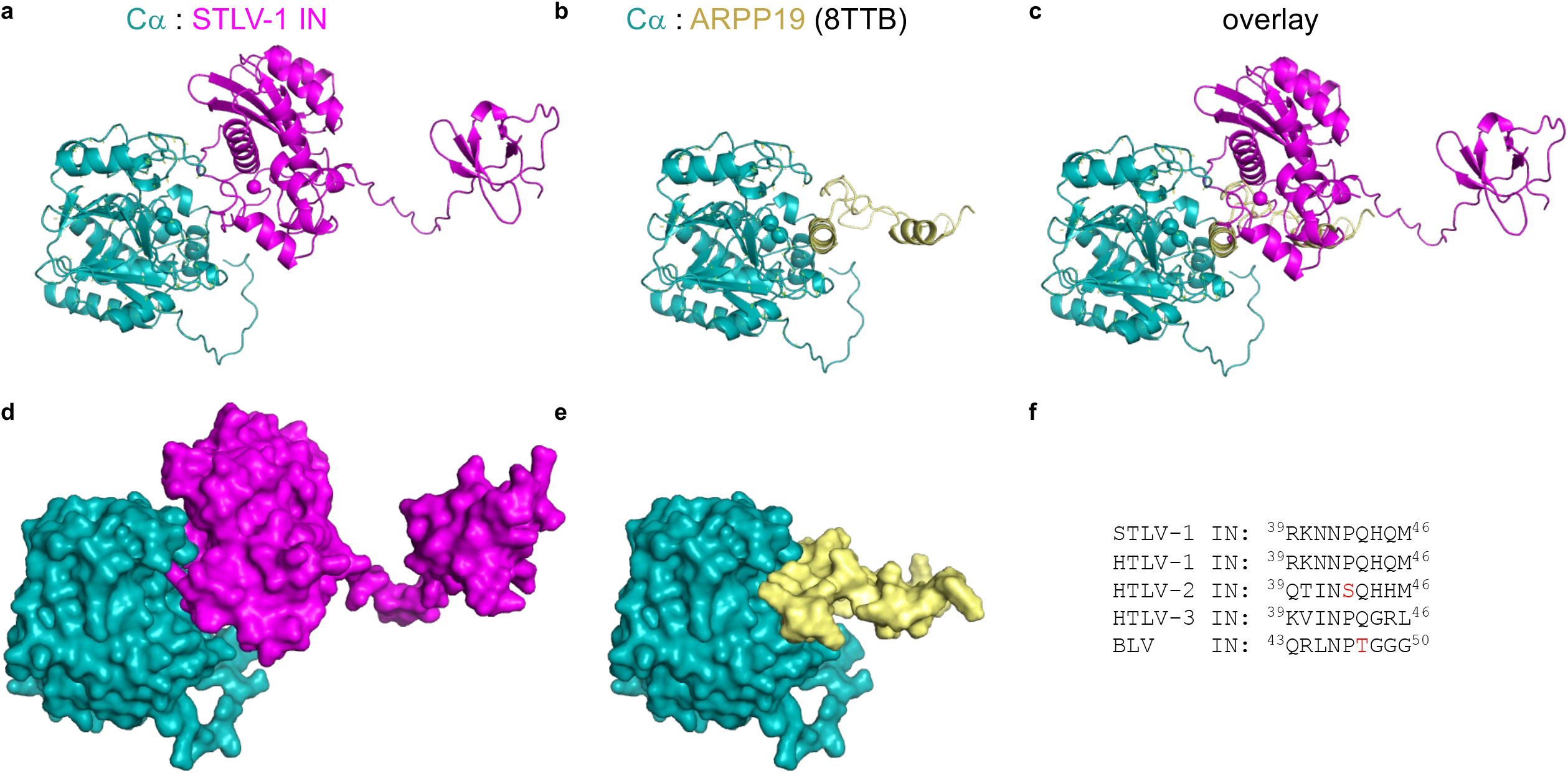
Deltaretroviral IN mimic endogenous PP2A-B55 inhibitor ARPP19 to block activity of PP2A-B56. ***a***, Cartoon representation of Cα bound to the outer IN molecule. ***b***, cartoon representation of Cα bound to inhibitor ARPP19 (PDB ID: 8TTB)[35], ***c***, overlay of ***a*** and ***b. d***, and ***e*** are surface representations of panels ***a*** and ***b*** respectively. ***f***, Alignment of STLV-1 IN residues ^39^KNNPQHQM^46^ with homologous residues in other deltaretroviral INs.

Free deltaretroviral INs can bind to each of the five members of the human B56 family, and in the context of the intasome, B56 residues critical for the interaction are highly conserved (Supplementary Figure S8). Budding yeast expresses a single B56 orthologue. Gene duplication over time resulted in higher eukaryotes to express five orthologues, B56α-ε with partially redundant roles. The question remains whether all five B56 isoforms (can) play a role in HTLV-1 infection. Bhatt *et al*. suggested B56γ, but not B56α plays a role in establishing HTLV-1 infection [13]. Further work is needed to dissect the contribution of the different B56 isoforms and to uncover the mechanism by which PP2A-B56 supports HTLV-1 infection. A major limitation of the HTLV-1 infection assays is their very low infection efficiency (~1% for the single-round WT vector and ~0.1% for the WT molecular clone), which has prevented us from sequencing enough unique integration sites under conditions lacking B56 binding. This makes it difficult to assess whether PP2A-B56 influences integration site selection. Crispr/Cas9 mediated knock-out of individual B56 isoforms could illuminate which of the five major isoforms plays a role.

PP2A-B56 is a highly abundant enzyme that can make up to 0.1-1% of the total cellular protein [37]. In 2017, Bekker-Jensen *et al*. determined an essentially complete HeLa proteome [38]. Whilst there are differences between certain cell lines, e.g. the HEK293 cell proteome expression profile is more akin to neuroblastoma cells, the overall expression profile is very comparable. In this work the authors were able to determine an approximate copy number of each of the detected cellular proteins. Notably, of the PP2A B56 regulatory subunits, B56δ is the most abundant with approximately 398,000 copies per HeLa cell [38]. Although this estimate is derived from HeLa cells rather than T cells, it provides an order-of-magnitude proxy for PP2A-B56 abundance. Given that a retroviral particle contains 50-100 IN molecules, it is highly unlikely that the function of IN would be to block cellular PP2A-B56 activity. Since IN is not a substrate of PP2A-B56, and PP2A inhibitors do not block HTLV-1 infection, our data suggests that the interaction of PP2A-B56 with the deltaretroviral intasome serves a structural and/or chromatin tethering role.

In conclusion, deltaretroviruses, such as the human pathogen HTLV-1, have evolved to use molecular mimicry to associate with the abundant Ser/Thr phosphatase PP2A-B56. In addition to the SLiM motif, IN also functionally mimics an endogenous PP2A-B55 inhibitor, ARPP19, and blocks access to the active site of Cα. We show here that the interaction between the HTLV-1 integration machinery and PP2A-B56 is essential to establish infection, while the PP2A-B56 phosphatase activity is not. Future work will be needed to elucidate the molecular mechanism by which PP2A-B56 supports HTLV-1 infection and integration.

## Materials and Methods

### DNA constructs

pET28aSUMO-LEDGF(1-324)-PPP2R5C2 was generated by PCR amplification of the PPP2R5C2 open reading frame (ORF) using primers GNM816 and GNM367 (Table S3) and pQFlag-PPP2R5C2 [11], and replacing the PPP2R5C(11-380) fragment from pET28aSUMO-LEDGF(1-324)-PPP2R5C(11-380) [12] using EcoRI and SalI restriction enzymes. HTLV-1 IN was expressed from pET28a-SUMO-H6P-HTLV-1 IN^s^ [12]. HTLV-1 IN^s^ mutants D122N, L213A, I216A, E218A and L213A/I216A were generated by site directed mutagenesis. B56γ(11-380) was expressed from pET28aSUMO-PPP2R5C(11-380) as described previously [12]; mutants I227A, I231A, H187A, and Y190A were generated by site directed mutagenesis. All primers used are described in Supplementary Table S3. pET28aSUMO-PPP2R5C(11-380) R188A and R197A to express B56γ(11-380)/R188A and B56γ(11-380)/R197A were previously described [11]. All plasmids were verified by Sanger sequencing. Baculoviral vector pOET1-His_6_-TEV-HA-PPP2CA was generated using *flash*BAC ULTRA™ by Oxford Expression Technologies, to express His_6_-TEV-HA-PPP2CA in insect cells.

pQ-HTLV-1 IN^s^-Flag-IRES-puroR to express HTLV-1 IN with a C-terminal Flag tag in human cells was described previously [11]. Mutations L213A, I216A, E218A and L213A/I216A were introduced using site directed mutagenesis. pQHA.2-PPP2R5E-IRES-hygroR was generated by cloning the PPP2R5E open reading frame into pQHA.2-IRES-hygroR in between MfeI/XhoI restriction sites.

### Recombinant protein expression and purification

The catalytic PP2A subunit Cα was produced in insect cells as His_6_-TEV-HA-Cα fusion protein. Hereto, *Trichoplusia ni* Tn5 cells grown in a shaking incubator (110 rpm) at 28°C in ESF-921 medium (Oxford Expression Technologies) to a density of 1×10^6^ cells/ml and infected with FBU-His_6_-TEV-HA-PPP2CA P2 virus at an MOI of 5. 72 h post-infection, the cells were harvested, and pellets frozen at −80°C until further processing. Pellets were thawed at room temperature and moved to ice immediately following thawing. All further steps were done on ice or at 4°C. Pellets were resuspended in ice cold lysis buffer 25 mM Tris-HCl pH 8.0, 100 mM NaCl, 5 mM DTT, supplemented with Complete EDTA free (Roche). Triton X-100 was added to a final concentration of 1% and the suspension was kept on ice for 10 min to allow lysis to occur. The suspension was sonicated (Branson sonifier, 40 sec, 70%, output 5) followed by centrifugation at 20 000 *g* for 1 h at 4°C. The supernatant was bound to Ni^2+^ resin (Cytiva) for 30 min at 4°C in the presence of 20 mM imidazole. The column was washed in lysis buffer supplemented with 20 mM imidazole. The protein was eluted by increasing imidazole concentration to 200 mM. One ml fractions were collected, analysed by SDS-PAGE and positive fractions pooled. The His_6_-tag was removed by TEV-protease cleavage overnight (at a ratio of 50:1 Cα:TEV) at 4°C. Next day, Cα was further purified by anion exchange chromatography. Briefly, the column was equilibrated with 25 mM Tris-HCl pH 8, the sample was loaded at 50 mM NaCl, the column extensively washed in 25 mM Tris-HCl pH 8.0, 50 mM NaCl and the protein was eluted by a linear 0.05 – 1 M NaCl gradient. Fractions were analysed by SDS-PAGE, and the protein product was further purified by size exclusion chromatography through a Superdex 200 column in 25 mM Tris-HCl pH 8.0, 100 mM NaCl. Positive fractions analysed by SDS-PAGE were pooled, supplemented with 2 mM DTT concentrated using Vivaspin with a molecular weight cut-off of 10 kDa to ~0.2 mg/ml, supplemented with 10% glycerol, aliquoted and flash frozen in liquid nitrogen.

STLV-1 and HTLV-1 IN proteins, LEDGF(1-324)-B56γ2 full-length (FL), further referred to as ΔIBD-B56γ, the scaffold PP2A subunit Aα, regulatory PP2A subunit B56γ(11-380) and their described mutant forms were expressed in *Escherichia coli* and purified as described previously [12, 32] [11]. Note, STLV-1 and HTLV-1 IN proteins were expressed as His_6_-SUMO-H6P-fusion. Thus, for His_6_-tagged versions of HTLV-1 IN, the His_6_-SUMO peptide was removed using ULP1 cleavage following affinity purification; to produce untagged HTLV-1 IN, the His_6_-SUMO-H6P-tag was removed by incubating the affinity purified protein with HRV 3C protease overnight before further purification by ion exchange and size exclusion chromatography [12].

To assemble and purify PP2A-core Aα and Cα were mixed in a 1: 2 molar ratio in 25 mM Tris pH 8.0, 100 mM NaCl and incubated on ice. The complex was then isolated by size exclusion chromatography. Fractions containing the PP2A core were collected and snap frozen in liquid nitrogen and stored at −80°C until further use.

### *In vitro* integrase strand transfer assay

The donor vDNA that mimics the 3’-processed LTR of HTLV-1 DNA was preassembled by annealing the U5S20Q-UP and U5S20Q-B oligonucleotides (Supplementary Table S3) in a reaction buffer containing 100 mM Tris-HCl pH 7.4 and 400 mM NaCl. The strand transfer reaction was set up as previously described by [32]. Briefly, in a final volume of 145 μL the following components were added: 24.3 mM 1,4-piperazinediethanesulfonic acid (PIPES) pH 6.0, 115 mM NaCl, 16.6 mM MgCl_2_, 5.8 µM ZnCl_2_, 8.73 mM DTT, 0.53 µM donor DNA, and 4 μg WT or mutant HTLV-1 IN. The reaction was incubated for 15 min at RT. After this time, 5 µL pGEM-9Zf(-) supercoiled plasmid DNA (60 ng/μL) was added and the reaction was incubated for 1 h at 37°C. To stop the reaction 0.5% SDS and 25 mM EDTA pH 8.0 were added. Protein degradation was obtained by incubating the reaction for 1 h at 37°C with 30 μg proteinase K (Roche). The DNA was precipitated overnight at −20°C with 70% (v/v) ethanol in the presence of 20 μg glycogen. The pellet was resuspended in 1.5X agarose loading dye and the DNA products were finally separated by electrophoresis through 1.5% agarose gel, and detected by staining with ethidium bromide. The ImageLab 4.1 software (Bio-Rad) was used to quantify the concerted integration products.

To investigate stimulation of HTLV-1 IN activity by B56, strand transfer assays were set-up as above with some minor modifications. The strand transfer reaction was prepared on ice. The components were added in the following order in a total volume of 20 μL: 16 μL reaction buffer (223 mM HEPES pH 7.1, 18.7 mM MgCl_2_, 25 mM DTT, 10.9 μM ZnCl_2_), 2 μL donor vDNA (40 μM), 1 μL of B56γ(11–380) (0-30 μM), and 1 μL of 60 μM WT or mutant IN. Different concentrations of B56 were used and are indicated in the corresponding figures. The reaction was incubated for 10 min at RT and 10 μL of 30 ng/μL pGEM-9Zf(-) supercoiled plasmid DNA was added. The reaction was incubated for 90 min at 37°C and stopped by adding 0.5% SDS/25 mM EDTA, pH 8.0. Proteins were degraded by adding 30 μg of proteinase K (Roche) for 1 h at 37°C. The DNA was precipitated overnight at −20°C with 70% (v/v) ethanol supplemented with 20 μg glycogen and the pellet was resuspended in 1.5X agarose loading dye. The DNA products were separated by electrophoresis (1.5% agarose gel stained with ethidium bromide) and the DNA product that corresponds to the concerted integration was quantified by densitometry using the ImageLab 4.1 software (Bio-Rad). Data was analysed in Prism v10.

### Electrophoretic mobility shift assays

STLV-1 IN(A219E) was mixed on ice with either ΔIBD-B56γ FL or with PP2A-B56γ FL holoenzyme and left at 4°C for minimally 30 min before proceeding. Complexes were assembled at a 1:1 molar ratio and a final concentration of 5 μM in 25 mM Tris pH 7.4, 200 mM NaCl, 2 mM DTT. Intasome assemblies took place at a final concentration of 1 μM IN : PP2A-B56 with 1 μM Atto-680 labelled vDNA mimics [12] in a buffer containing 20 mM BTP pH 6.0, 10 mM CaCl_2_, 10 mM DTT, 10 μM ZnCl_2_ were added in reaction buffer and incubated at 37°C for 10 min. The complex was then moved to room temperature, and NaCl was added to a final concentration of 1.2 M. Heparin was added to a final concentration of 10 μg/ml and the samples were incubated at room temperature for different lengths of time. 6 μl sample was separated on a 3% low melting point agarose gel supplemented with 10 μg/ml heparin and run in 1x TBE buffer at 120V, 400 mA for 90 min. Gels were imaged using the Azure c600.

### Phosphatase assays

PP2A core was assembled by mixing purified Cα and Aα in 25 mM Tris HCl pH 8, 100 mM NaCl at a final concentration of 250 nM. The colorimetric malachite green phosphatase assay was used to measure PP2A enzymatic activity using the PP2A specific phospho-Threonine peptide (K-R-pT-I-R-R) as a substrate. Absorbance was measured at 620nm. A standard curve was made using a dilution series of potassium phosphate ranging from 0 to 2000 pmoles phosphate. Reactions with the phospho-Thr substrate were done in the following phosphatase assay buffer: 25 mM Tris-HCl pH 7.4, 1 mM EDTA, 1 mM EGTA, 1 mM DTT and 0.25 mg/ml BSA and allowed to take place for 20 min at 37°C. Each reaction contained 20 nM PP2A-core, and 80 μM phosphopeptide. For each assay, a positive control was included which contained the PP2A core mixed with the dilution buffer the IN or the intasome preparation was made in, and a negative control which contained PP2A core in the presence of 1x PhosSTOP (Sigma). All buffers were tested as blank. For inhibition by the HTLV-1 intasome, pre-assembled and purified PP2A-core was used. PP2A core and IN or intasome were pre-incubated on ice for 10 min before addition of the phosphopeptide. Reactions were then allowed to take place at 37°C for 20 min before the malachite green substrate was added. Absorbance was measured following a 15-min incubation at room temperature with the colorimetric substrate. Data shown are from three biological replicates (free IN), or two biological replicates (intasome assemblies). Note that WT or SLiM mutant STLV-1 IN and WT HTLV-1 IN were used for these assays as indicated in the figure. Each biological replicate represent the average of three technical replicates. Graphs were generated in Prism v10.

### Fluorescence polarization assays

FP assays were conducted in low protein binding, black 384-well plates (Corning) by measurement of the perpendicular and parallel fluorescence signals at 520 nm (the emission wavelength of fluorescein) from each well. Samples were prepared with decreasing concentrations of B56γ(11-380) variants (between 32 – 0.125 µM) in 25 mM Tris pH 8.0, 300 mM NaCl supplemented with 40 nM of the appropriate peptide. The following peptides were used: HTLV-1 IN P1: FAM-Ahx-KTRWQLHHSPRLQPIPETHSLS-Amide; and RepoMan: FAM-Ahx-GERDIASKK PLLSPIPELPEVP-Amide. Polarization values were calculated according to the following equation: mP =(I_parallel_ – I_perpendicular_)/(I_parallel_ + I_perpendicular_). The resulting mP values corresponding to each well were plotted against the protein concentration, and a curve was fitted for anisotropic binding in Prism v8 to determine the equilibrium constant for each peptide/protein pair.

### His_6_-tag pull-down, Flag-immunoprecipitation and western blot detection

His_6_-tag pull-downs were done as described in [11]. Flag-immunoprecipitation was done as described in [11] using PP2A IP buffer (25 mM Tris-HCl pH 8.0, 0.1% Nonidet-P40, 150 mM NaCl, 3 mM EDTA, 3 mM EGTA supplemented with Complete EDTA free (Roche)). Following separation on 11% SDS-PAGE the proteins were transferred to nitrocellular membrane using wet transfer. Blots were blocked in blocking buffer (5% milk in phosphate buffered saline (PBS)). Primary and secondary antibodies were diluted in blocking buffer supplemented with 0.1% Tween-20. In between antibody incubations the blots were washed extensively with PBS-Tween (PBS supplemented with 0.1% Tween-20). Detection was done using Clarity or ClarityMax (BioRad) and developed using a c600 Azure imager, and signals were quantified using the Azure software package. A list of antibodies used is available in Supplementary Table S4.

### HTLV-1 co-culture infection assays

Single round HTLV-1 infection assays [24] were done using the method described in [13]. To generate infectious virus from the pACH molecular clone, 293T cells were transfected by pACH complexed with XtremeGene-HP. Cell cultures were incubated for 16 h before cells were washed and treated with DNase I (Sigma). Cell culture media was replaced, and 24 h later, cells were harvested and gamma-irradiated (40,000 Rad). For infection assays into HOS reporter target cells, irradiated cells were co-cultured with HOS cells at a 5:1 producer: target cell ratio in DMEM lacking serum and supplemented with 15 mM MgCl_2_ for 16 h at 37°C, in 5% CO_2_ atmosphere. Following co-culture, puromycin-resistant HOS target cells were passaged 5 times in the presence of puromycin for 12 days.

For infection via MT-2 cells, Jurkat target cells were treated with PP2A inhibitors 4 h prior to infection, or raltegravir 24 h prior to infection. Target cells were co-cultured with irradiated MT-2 cells at a 1:1 ratio in serum-free RPMI. After 24 h, MT-2 cells were depleted using anti-CD25 magnetic dynabeads as per the manufacturer’s instructions. The resultant culture was passaged over 16 days in the presence of appropriate cell culture additives.

### Western Blot detection of HTLV-1 proteins in producer cells

To prepare inactivated lysates from producer cell lines, cells were lysed on ice in buffer supplemented with 0.4% NP-40 before storage at −150°C. Following this, samples were briefly thawed and separated by centrifugation for 20 minutes at 17,000 *g* at 4°C and supernatants were separated by SDS-PAGE and western blotting. In the case of assaying Tax expression immunoprecipitation was performed on the lysate using mouse anti-Tax antibody (sc-57872, Insight Biotechnology) bound to protein A agarose beads. Elution yielded a 25-fold enrichment in Tax protein compared to the input lysate that was clearly detectable by western blotting. For detection of soluble virion-associated p24^Gag^, culture supernatants were ultracentrifuged for 2 h at 87,500 *g* through a 20% sucrose-cushion. Supernatant was removed and resultant pellets were resuspended in ice cold PBS for analysis by SG-PERT and western blot analysis. Equal volumes of samples were separated by SDS-PAGE and transferred to nitrocellulose membrane for western blot detection as described above.

### Statistical analyses

Prism v10 (GraphPad) was used for statistical tests and graphs in Figures 1-3 and 6. One-way ANOVA was used to compare values in experimental groups to those of control groups, with Dunnet’s post-hoc test applied. P values for comparisons are represented by asterices in the figures; \**p* < 0.05, ** *p* < 0.01, *** *p* < 0.001, **** *p* < 0.0001.

### Cryo-EM grid preparation and data collection

The intasome was assembled using recombinant STLV-1 IN(A219E), ΔIBD-B56γ full-length chimera, synthetic oligonucleotides, and purified by size exclusion chromatography as described [12, 32]. The peak fraction was incubated with 50 μM bictegravir and 20 mM MgSO_4_ at room temperature for 10 min. Next, the intasome was supplemented with 1.5-molar excess of Aα (~18 μg/ml) and Cα (~9 μg/ml) and an equal volume of 25 mM Tris-HCl pH 7.4 to reduce the NaCl concentration to 150 mM NaCl. The holoenzyme was allowed to form at 4 °C for 15 min. UltraAuFoil R 1.2/1.3 Au 300-mesh grids (Electron Microscopy Sciences) [39] were glow-discharged for 4 min at 45 mA in an Emitech K100X instrument (Electron Microscopy Sciences) and covered with a layer of graphene oxide (Sigma-Aldrich, catalogue #763705) [40] for immediate use. 4 μL intasome was spotted on the graphene side of a coated grid. The grids were incubated for 1 min at 22 °C and 95 % humidity, blotted for 2.5 s prior to plunge-freezing into liquid ethane using a Vitrobot Mark IV instrument (Thermo Fisher Scientific).

Single particle images were acquired on a 300-keV Titan Krios cryo-electron microscope (Thermo Fisher Scientific) with a Gatan GIF BioQuantum energy filter and a K3 Summit direct electron detector (Gatan). Micrographs were recorded in dose-fractionation mode, at a calibrated magnification corresponding to 0.85 Å per physical pixel (0.425 Å per super-resolution pixel) at the detector. A total electron exposure of 55.2 e/Å^2^ (corresponding to 40 e/pixel^2^) was fractionated across 40 movie frames. An 20-eV energy slit and a defocus range of −1.4 to −3.6 μm were used to acquire a total of 31,146 micrograph movie stacks.

### Image processing and 3D reconstruction

Micrograph movie frames were aligned, binned to the physical pixel size, and summed with dose weighting using MotionCor2 v1.5.0 [41]. Contrast transfer function (CTF) parameters were estimated using Gctf-v1.18 [42]. Micrographs with crystalline ice contamination were discarded at this stage, leaving 27,970 images for further processing (Supplementary Fig. S5a). An initial subset of particles, picked using crYOLO [43] with the general model, was extracted with 2x binning. These particles were subjected to several rounds of 2D classification in CryoSPARC v4.1 [44], and 51,026 particles belonging to well-defined classes were used to train Topaz [45] for automated picking across the entire dataset. A total of 1,641,876 particles were subsequently extracted with 4x binning (final box size of 146 pixels) and subjected to iterative rounds of 2D classification in CryoSPARC v4.1 [44] and Relion-4.0 [46]. This procedure yielded 411,539 particles contributing to well-defined 2D classes (Supplementary Fig. S5a). These particles were re-extracted with 2x binning (final box size of 292 pixels) and subjected to 3D classification into two classes using ab initio reconstruction followed by heterogenous refinement in CryoSPARC v4.1 without imposing symmetry. A subset of 244,887 particles from the best-resolved class was re-extracted at a pixel size of 1.133 Å with a box size of 480 pixels (corresponding to a binning factor of 4/3) and used for 3D reconstruction in CryoSPARC v4.7.1. Following Bayesian particle polishing in RELION-5.0 and per-particle CTF refinement in CryoSPARC v4.7.1, the final global reconstruction was obtained using non-uniform refinement in CryoSPARC v4.7.1 with *C2* symmetry imposed, yielding an overall resolution of 2.80 Å (Supplementary Fig. S5b). In addition, a local reconstruction was performed to improve the resolution of the PP2A scaffold subunit Aα and to facilitate model building. To this end, a symmetry-expanded particle set was subjected to local refinement in CryoSPARC v4.7.1 using a mask focused on one half of the particle. Refinement was performed with per-particle scale minimization enabled, while all other parameters were kept at their default values, yielding a reconstruction with an overall resolution of 2.63 Å within the masked region. The particles were subsequently subjected to 3D classification into three classes without alignment in RELION-4.0, using the same local mask. Classification was run for 45 iterations with a regularisation parameter (T) of 8. This procedure isolated a single well-defined class comprising 338,454 particles. These particles were then used for a final round of local refinement in CryoSPARC v4.7.1, resulting in a reconstruction of the masked half at an overall resolution of 2.59 Å (Supplementary Fig. S5c).

Cryo-EM map resolutions were determined using the gold-standard Fourier shell correlation (FSC) 0.143 criterion [47, 48] (Supplementary Table S2); local resolutions were estimated in CryoSPARC-4.7.1 (Supplementary Figure S5b,c). To aid in model building process and to prepare figures, the maps were filtered and sharpened using EMReady v2 [49]. For real-space refinements of the atomistic models, the cryo-EM reconstructions were sharpened and locally filtered based on local resolution estimations using CryoSPARC-4.7.1.

### Model building and refinement

Model building was initiated using the structure of the STLV-1 intasome in complex with B56 and bictegravir (PDB accession code: 7OUH) [32]. The IN A219E mutation was introduced, and local conformational adjustments of selected residues and viral DNA bases were performed in Coot [50]. The PP2A holoenzyme components were modelled starting from the PP2A–B56 complex structure (PDB accession code: 2NPP) [51]. Initial fitting of all components into the cryo-EM density maps was carried out in UCSF Chimera [52]. The combined model was then extended and manually adjusted in Coot using both global and locally refined cryo-EM reconstructions. The full atomistic model was refined by real-space refinement in Phenix (v2.0-5936) [53] against the locally filtered cryo-EM map. Refinement included secondary structure restraints, base-pairing and base-stacking restraints derived from the model, as well as metal coordination restraints. Non-crystallographic symmetry (NCS) restraints were applied to the two halves of the symmetric nucleoprotein assembly. The quality of the final model was assessed using MolProbity [54] (Supplementary Table S2).

### Negative stain grid preparation and data collection of HTLV-1 intasome

The HTLV-1 strand transfer complex intasome was assembled using HTLV-1 IN D122N (active site mutant) under conditions described above for the STLV-1 intasome, using oligonucleotides listed in Supplementary Table S3 [13]. No strand transfer inhibitor was added to the purified assemblies before grid preparation and data collection. 300 mesh copper grids with continuous carbon (EM resolutions) were glow discharged using Emitech 1 min 45 mA – 3 μl HTLV intasome:B56 or intasome:PP2A-B56 were applied to freshly glow discharged carbon – followed by negative staining with 2% uranyl acetate.

## Data availability

The global and local cryo-EM reconstructions will be deposited with the EMDB once the manuscript has been accepted for publication. Refined coordinates of the complete assembly will be available with PDB upon acceptance of publication (Supplementary Table S2). Raw cryo-EM data ara vailable upon request. The authors declare that all other data supporting the findings of this study are available within the paper, and it supplementary files. Correspondence and requests for materials should be addressed to G.N.M.

## Supplementary data

Supplementary data are available on *Nat Comms* online.

## Acknowledgements

We are grateful to Chou-Zhen Giam for sharing the HOS reporter cell line with us [28].

## Author contributions

JJM: Investigation, formal analysis, validation, visualization, writing – review and editing. EP: investigation, formal analysis, writing – review and editing. TV: investigation, formal analysis, validation, writing – review and editing. RS: investigation, formal analysis, validation, visualization, writing – review and editing. NC: investigation. PC: formal analysis, validation, visualization, funding acquisition, writing – review and editing. GNM: conceptualization, project administration, investigation, formal analysis, validation, visualization, funding acquisition, supervision, writing – original draft.

## Supplementary Information

**Supplementary Table S1** | **Quantification of HTLV-1 IN and RepoMan binding to B56γ(11-380)**. WT or mutant B56γ(11-380) were used to determine binding affinity to WT HTLV-1 IN peptide and RepoMan peptide.

**Supplementary Table S2** | **Cryo-EM data**.

**Supplementary Table S3** | **Primer sequences used in this paper**.

**Supplementary Table S4** | **Primary and secondary antibodies used in this manuscript**.

**Supplementary Figure S1**| **Fluorescence polarization experiments with B56γ(11-380) WT and mutants, and HTLV-1 peptides. *a***, Peptides used in the fluorescence polarization assay. The LxxIxE residues are highlighted in red. ***b***, surface representation of B56γ(11-380) bound to the HTLV-1 SLiM peptide (PDB ID:6TOQ)[12]. ***c***, Fluorescence polarization data using WT HTLV-1 IN peptide and WT or mutant B56γ(11-380). ***d***, Fluorescence polarization curves using RepoMan peptide and WT or mutant B56γ(11-380).

**Supplementary Figure S2** | **HTLV-1 IN SLiM mutants binding to all 5 B56 isoforms. *a***, Representative western blot showing Flag-immunoprecipitation of WT IN, IN(D122N), IN(L213A), IN(I216A), IN(E218A) and IN(L213A/I216A) and the recovery of endogenous B56α, β, γ, and δ. β-actin was used as loading control. As expected, HTLV-1 IN does not bind to striatin, B55α or PR72 [11]. ***b***, Flag-immunoprecipitation as in ***a***, using Flag-tagged WT and mutant HTLV-1 IN proteins as bait, and HA-tagged B56ε as prey. Antibodies are shown on the right. Mutants are indicated on the top of the western blots.

**Supplementary Figure S3** | **HTLV-1 infection assay using the pACH molecular clone and a GFP-reporter cell line. *a***, Schematic illustrating the set-up of the co-culture experiment made using BioRender. ***b***, SG-PERT assay measuring RT activity in the supernatant of the producer cells. ***c***, Time-course of GFP expression for pACH^WT^ (WT), pACH^ΔENV^ (ΔENV) and pACH^IN(D122N)^.

**Supplementary Figure S4** | **Characterization of STLV-1 intasome : PP2A-B56**γ **holoenzyme complexes by EMSA and negative stain EM. *a***, electrophoretic mobility shift assay (EMSA) using Atto680-labeled viral DNA mimics, STLV-1 IN, B56 γ full-length, and PP2A-B56γ (Aα:Cα:B56γ) holoenzyme. Following assembly and resuspension in high salt, the complexes were incubated on ice for 5 min, 1h or 2h as indicated above the gels. Migration of the different nucleoprotein complexes are indicated on the right. *Lanes 1:* negative control (vDNA only), *lanes 2*: STLV-1 intasome – B56γ; *lanes 3*: STLV-1 intasome – PP2A-B56γ. ***b***, 2D class averages of STLV-1 intasome – B56γ (top) and STLV-1 intasome – PP2A-B56γ (bottom) as indicated above the panels. Scale bars are 10 nm. ***c***, average length of the 2D negative stain EM particles.

**Supplementary Figure S5** | **Cryo-EM analysis of STLV-1 intasome in complex with PP2A-B56γ holoenzyme. *a***, Representative cryo-EM micrograph showing a subset of STLV-1 intasome particles (*left*, circled) and corresponding 2D class averages (*right*). Scale bars, 20 nm. ***b***, *Top*: Gold-standard half-map Fourier shell correlation (FSC) curves for the full reconstruction without masking (blue), with loose mask (green), a tight maks (ref), and after mask correction (purple). The FSC = 0.143 criterion (black line) corresponds to an overall resolution of 2.80 Å. *Bottom left:* Angular distribution of particle orientations contributing to the final reconstruction. *Right:* Orthogonal slices through the density map coloured by local resolution, ranging from 2.5 Å (blue; core regions including IN and B56γ) to lower resolution (red) at the periphery of the scaffold subunit. ***c***, Half-map FSC curves, viewing angle distribution, and local resolution map for the local reconstruction of one half of the complex.

**Supplementary Figure S6** | **Examples of cryo-EM map for select regions of the structure. *a***, Left: PP2A Cα subunit (chain E) is shown in stick representation (carbon atoms in cyan), overlaid with the IN^39-60^ loop (chain A residues 38-53 are shown as a magenta cartoon). The cryo-EM map from local refinement, filtered with EMReady, is shown as a semi-transparent grey surface. The location of the Cα active site is indicated by a dashed circle. *Right:* Extent of cryo-EM map coverage for the IN loop. The loop is shown as a cartoon with side chains supported by the density displayed as sticks; PP2A Cα is shown as a cartoon with active-site residues in sticks and the metal atoms (modelled in the structure as Mg^2+^ ions) as grey spheres. IN residues 44-49 are not resolved in the map. ***b***, Cryo-EM density map for the PP2A Aα scaffold subunit is shown as a semi-transparent grey surface, with the atomic model in stick representation (carbon atoms in grey). Selected residues are labelled where possible without obscuring the view of the map. ***c***, Cryo-EM map of the inner IN subunit (chain B), including the active site bound to vDNA and the strand transfer inhibitor bictegravir (BIC). The protein, vDNA, and inhibitor are shown as sticks with carbon atoms coloured light pink, blue, and yellow, respectively. Grey spheres represent Mg^2+^ ions, and red spheres indicate coordinated water molecules.

**Supplementary Figure S7** | **PP2A-B56γ holoenzyme in intasome structure superimposed with free PP2A-B56γ**. PP2A-B56γ bound to the STLV-1 intasome (chain C: Aα; chain D: B56γ; chain E: Cα; this study) was superimposed with the previously solved PP2A-B56γ holoenzyme structure (PDB ID: 2NPP) [51]. Both structures are shown in cartoon representation, colours for each of the chains are indicated in the figure.

**Supplementary Figure S8** | **The STLV-1 IN interaction interface on B56γ is highly conserved across all B56 isoforms**. Alignment of human B56α-ε isoforms. The canonical SLiM binding site is indicated with a red box, and the novel identified interaction interface described in this work is marked with a green box. ^*^ mark conserved residues involved in direct interaction with IN, orange arrow indicates residue that is not 100% conserved across all 5 isoforms.

