## Supplementary Table S1 for "HTLV-1 intasome recruits the PP2A-B56 holoenzyme and restricts its phosphatase activity"

**Supplementary Table S1- Quantification of HTLV-1 IN and RepoMan SLiM binding to B56γ(11-380)**

| B56γ (11-380) | HTLV-1 IN (μM) | | | RepoMan (μM) | | |
|---|---|---|---|---|---|---|
| | KD | SD (+/-) | $R^2$ of fit | KD | SD (+/-) | $R^2$ of fit |
| WT | 1.323 | 0.08913 | 0.9959 | 1.886 | 0.2311 | 0.9842 |
| I227A | 7.779 | 1.661 | 0.9627 | 6.845 | 1.188 | 0.9735 |
| I231A | 5.674 | 0.5785 | 0.9899 | 9.266 | 4.636 | 0.837 |
| H187A | NB | NB | NB | NB | NB | NB |
| Y190A | 9.35 | 2.506 | 0.9474 | NB | NB | NB |
| R188A | 2.337 | 0.2057 | 0.9914 | 3.367 | 0.3238 | 0.9897 |
| R197A | 17.4 | 5.597 | 0.949 | NB | NB | NB |

NB: no binding observed

**Table S2.** Cryo-EM data collection, image processing, and model refinement.

| **Data collection** | **STLV intasome in complex with PP2A holoenzyme** | |
|---|---|---|
| Microscope, operating voltage | Titan Krios, 300 keV | |
| Detector | Gatan K3 | |
| Energy filter | Gatan GIF BioQuantum | |
| Energy selection slit (eV) | 20 | |
| Micrograph pixel size (Å) | 0.85 | |
| Underfocus range (nominal, µm) | 1.4 - 3.6 | |
| Number of frames per movie | 40 | |
| Total electron fluence (e/Å$^2$) | 55.2 | |
| Automation software | FEI EPU | |
| Number of micrograph movies/tilt series used | 27,960 | |
| **Reconstruction** | Local refinement | Global refinement (full structure) |
| **EMDB ID** | **XXXX** | **XXXX** |
| Software for 2D classification | cryoSPARC | cryoSPARC |
| Software for 3D classification | cryoSPARC, Relion | cryoSPARC, Relion |
| Software for final reconstruction | cryoSPARC | cryoSPARC |
| Number of refined particles | 338,454 | 244,887 |
| Pixel size for final reconstructions | | |
| Symmetry | *C1* | *C2* |
| Global resolution (FSC 0.143, Å) | 2.59 | 2.80 |
| Map resolution range (Å) | 2.5 - 5.5 | 2.5 - 6.5 |
| Map sharpening B factor | -76 | -83 |
| **Model refinement** | | |
| **PDB ID** | | **XXXX** |
| Software for real-space refinement | | Phenix |
| Model composition | | |
| Number of non-H atoms | | 31,134 |
| Number of protein residues | | 3,676 |
| Number of nucleotides | | 80 |
| Number of ligands (Zn, Mg, KLQ) | | 4, 8, 2 |
| *B* factors (Å$^2$) | | |
| Protein | | 67.4 |
| RNA | | 63.6 |
| Ligands | | 15.9 |
| Real-space correlation ($CC_{mask}$, $CC_{box}$, $CC_{peaks}$, $CC_{volume}$) | | 0.76, 0.71, 0.69, 0.74 |
| R.m.s. deviations | | |
| Bond lengths (Å) | | 0.003 |
| Bond angles (°) | | 0.525 |
| Model validation | | |
| MolProbity score | | 1.63 |
| Clash score | | 3.61 |
| Poor rotamers (%) | | 2.52 |
| CaBLAM outliers (%) | | 2.04 |
| Ramachandran plot quality (%) | | |
| Favored, Disallowed | | 97.0, 0 |

### Supplementary Table S3- Primer sequences

**Primers used for cloning**

| Primer name | sequence (5'-3') | description |
|---|---|---|
| GNM364 | ggcc caattg ATGTCCTCAGCACCAACTACTCCTC | sense PPP2R5E with MfeI site (for FL) |
| GNM365 | GGCCGTCGACTTAAGTTGGAATTATTCCATCACGTC | antisense PPP2R5E with stop codon and SalI site |
| GNM816 | ggccgaattcATGTTGACATGTAATAAAGCGGGC | sense PPP2R5C2 with EcoRI site |
| GNM367 | GGCCGTCGACCTAGCGGCCGTCCTGGG | antisense PPP2R5C2 with stop codon and SalI |
| GNM635 | GGCAGGGCCGTTGTTTGTGTTTATGTAGCTAGGCTTGCC | antisense HTLV-1 IN sequence to introduce D122N mutation |
| GNM636 | gctacataaacacaAacaacggccctgcctatatttcc | sense HTLV-1 IN D122N use together with GNM637 |
| GNM637 | TCTGCAATTGCTTACCCATGGTGTTGGTGGTCTTTTTCTTTGGGATC | antisense HTLV-1 IN sequence with MfeI site |
| GNM638 | CCCACGGTACCTATACGGTCCCGCTGGGCGC | sense HTLV-1 primer, anneals within RT, with KpnI site splice together using primer GNM637 |
| GNM711 | cccccgactccagccggccccagagacac | HTLV-1 IN I216A Forward (in pACH and pDS-601) |
| GNM712 | GTGTCTCTGGGGCCGGCTGGAGTCGGGGG | HTLV-1 IN I216A Reverse (in pACH and pDS-601) |
| GNM713 | ccccccgaGCCcagccgGCcccagagacac | HTLV-1 IN L213A/I216A Forward (in pDS-601) |
| GNM714 | GTGTCTCTGGGGCCGGCTGGGCTCGGGGGG | HTLV-1 IN L213A/I216A Reverse (in pDS-601) |
| GNM715 | ccccccgaGCccagccgatcccagagacac | HTLV-1 IN L213A Forward (in pACH and pDS-601) |
| GNM716 | GTGTCTCTGGGATCGGCTGGGCTCGGGGGG | HTLV-1 IN L213A Reverse (in pACH and pDS-601) |
| JM_86 | cacagccccagaGCgcagcccGCccccgagacacacag | Internal sense primer HTLV-1 IN L213A/I216A. A in Capitals. |
| JM_87 | ctgtgtgtctcggggGCgggctgcGCtctggggctgtggtg | Internal antisense primer for HTLV-1 IN L213A/I216A double mutant |
| RMG1 | ctgcaccacagccccagagcgcagcccatccccgagacac | HTLV-1 INs sense L213A |
| RMG2 | GTGTCTCGGGGATGGGCTGCGCTCTGGGGCTGTGGTGCAG | antisense L213A |
| RMG3 | cacagccccagactgcagccc gcccccgagacacacagcctgagc | HTLV-1 INs sense I216A |
| RMG4 | gctcaggctgtgtgtctcgggggcgggctgcagtctggggctgtg | antisense I216A |
| RMG5 | cccagctacatcaacacc aac aacggccctgcctatatcagc | HTLV-1 INs sense D122N |
| RMG6 | GCTGATATAGGCAGGGCCGTTGTTGGTGTTGATGTAGCTGGG | HTLV-1 INs antisense D122N |
| JM_105 | agccgatcccagCgacacattccctc | HTLV-1 IN E218A Forward (in pACH and pDS601) |
| JM_106 | gagggaatgtgtcGctgggatcggct | HTLV-1 IN E218A Reverse (in pACH and pDS601) |
| JM_76 | cagcccatccccgCgacacacagc | Internal sense for HTLV1 INs E218EA mutant |
| JM_77 | gctgtgtgtcGcggggatgggctg | Antisense HTLV1 INs E218A mutant |
| JM_66 | cttaaaaccacccttGCcagaatctatgggaaattcc | Internal Sense primer for PPP2R5C H187A |
| JM_67 | taggaatttcccatagattctgGCaagggtggttttaagaaaatc | Internal Antisense primer for PPP2R5C H187A |

| | | |
|---|---|---|
| JM_68 | acccttcacagaatcGCtgggaaattcctaggcttg | Internal Sense primer for PPP2R5C Y190A |
| JM_69 | gcctaggaatttcccaGCgattctgtgaagggtggttttaag | Internal Antisense primer for PPP2R5C Y190A |
| JM_70 | gcagagttactggaaatattgggaagtGCaattaatggatttgcc | Internal Sense primer for PPP2R5C I231A |
| JM_71 | ggcaaatccattaattGCacttcccaatatttccagtaactc | Internal Antisense primer for PPP2R5C I231A |
| JM_72 | gagttactggaaGCattgggaagtataattaatggatttgccttaccac | Internal Sense primer for PPP2R5C I227A |
| JM_73 | tatacttcccaatGCttccagtaactctgctatgccattatg | Internal Antisense primer for PPP2R5C I227A |

**Primers used for qPCR based assays**

| Primer name | Sequence (5' to 3') | Function |
|---|---|---|
| SK43 | CGGATACCCAGTCTACGTGT | qPCR for PVL (*tax* Forward) |
| SK44 | GAGCCGATAACGCGTCCATCG | qPCR for PVL (*tax* Reverse) |
| *gapdh* (forward) | AACAGCGACACCCATCCTC | qPCR for PVL |
| *gapdh* (reverse) | CATACCAGGAAATGAGCTTGACAA | qPCR for PVL |
| *SG-PERT* (forward) | TCCTGCTCAACTTCCTGTCGAG | SG-PERT |
| *SG-PERT* (reverse) | CACAGGTCAAACCTCCTAGGAATG | SG-PERT |

**Primers used to assemble HTLV-1 strand transfer complexes**

| Primer name | Sequence (5' to 3') |
|---|---|
| U5-25T20 | CCAGGAGAGAAATTTAGTACACAGATATCCACCCTAGTCAAGTGTGTCC |
| U5-nj25 | ACTGTGTACTAAATTTCTCTCCTGG |
| T20 | GGACACACTTGACTAGGGTG |

**Primers used for HTLV-1 strand transfer assays**

| | |
|---|---|
| U5S20Q-UP | GACTCACTATAGGGCACGCGTAGAGAAATTTAGTACACA |
| U5S20Q-B | ACTGTGTACTAAA TTTCTCTACGCGTGCCCTATAGTGAGTC |

**Supplementary Table S4- Antibody information**

| Antibody | Species | Company | Clone number | Catalogue number | Lot number |
|---|---|---|---|---|---|
| Anti-Flag | Mouse | Sigma | M2 | A8592-IM6 | SLBB9238 |
| Anti-HA tag | mouse | Sigma | 16B12 | H6908-.2ML | 97653 |
| Anti-Rat IgG-HRP | Rabbit | Abcam | Polyclonal | ab6734 | GR36157-10 |
| Anti-Rabbit IgG-HRP | Donkey | GE Life Sciences | Polyclonal | GE-NA934 | 6969611 |
| Anti-Mouse-IgG-HRP | Sheep | GE Life Sciences | Polyclonal | GE-NA931 | 14263051 |
| Anti-b-actin-HRP | Mouse | Abcam | AC-15 | ab49900 | GR151100-18 |
| Anti-B55a | Mouse | CST | 269.Ms | 5689T | 1 |
| Anti-B56a | Mouse | Insight Biotechnology | F-10 | SC-271151 | KO316 |
| Anti-B56b | Mouse | Insight Biotechnology | E-6 | SC-515676 | F1616 |
| Anti-B56g | Mouse | Insight Biotechnology | E-6 | SC-374380 | C1618 |
| Anti-B56d | Mouse | Insight Biotechnology | H5D12 | SC-81605 | B2117 |
| Anti-B56d | Rabbit | Bethyl Laboratories, Inc. | Polyclonal | A301-098A | A301-098A1 |
| Anti-B56e | Mouse | Insight Biotechnology | A-1 | SC-376176 | K1213 |
| Anti-PR72 | Rabbit | Bethyl Laboratories, Inc. | Polyclonal | A300-967A | A300-967A-T |
| Anti-STRIATIN3 | Rabbit | Bethyl Laboratories, Inc. | Polyclonal | AA304537A | AA304537A-T |
| Anti-HTLV-1 p19 | Mouse | Insight Biotechnology | TD-7 | SC-57870 | 4118 |
| Anti-HTLV-1 p24 | Rabbit | Abcam | 46/3.24.4 | ab9081 | 4 |
| Anti-HTLV-1 Tax | Mouse | Insight Biotechnology | IA3 | SC-57892 | F2817 |
| Anti-HTLV-1 gp46 | Mouse | Insight Biotechnology | 65/6C2.2.34 | SC-57865 | A0611 |
