## Supplementary Figures 1-8 for "HTLV-1 intasome recruits the PP2A-B56 holoenzyme and restricts its phosphatase activity"

### Supplementary Figure S1_Minnell

**a**

**HTLV-1 IN P1** 5’FAM-Ahx-KTRWQLHHSPRLQPIPETHSLS-Amide

**RepoMan** 5’FAM-Ahx-GERDIASKKPLLSPIPELPEVP-Amide

**b**

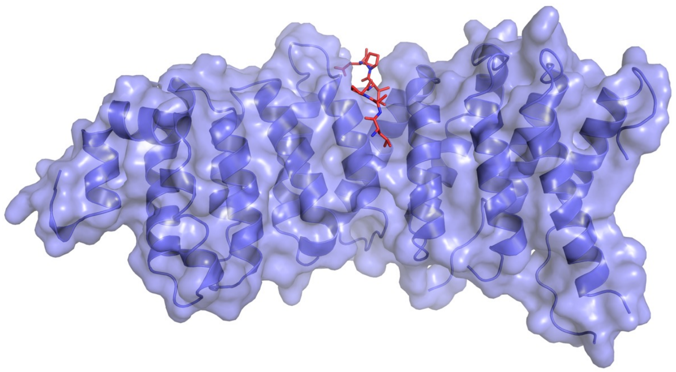

**c**

Binding of B56γ(11-380) to HTLV-1 IN peptide

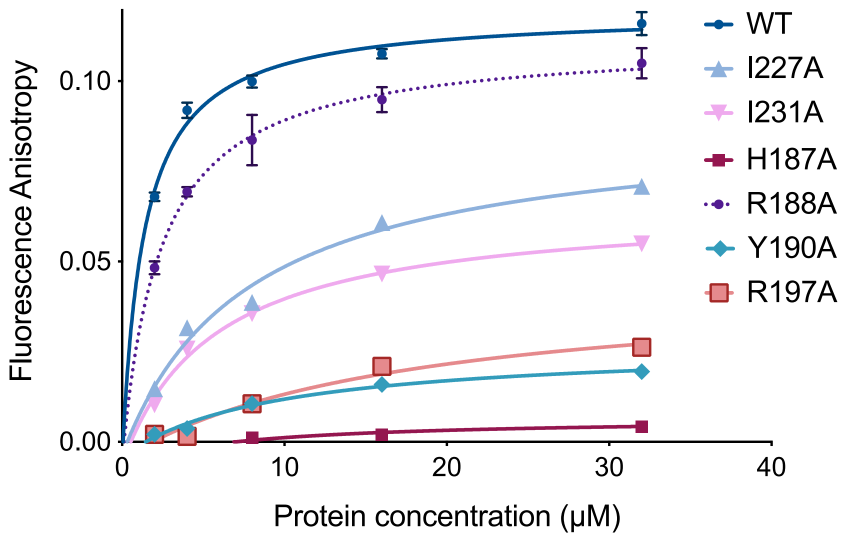


**d**

Binding of B56γ(11-380) to RepMan peptide

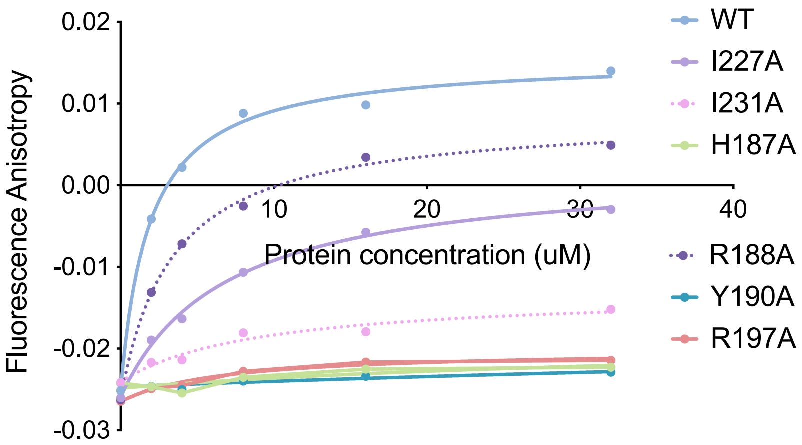

#### Supplementary Figure S2_Minnell

a

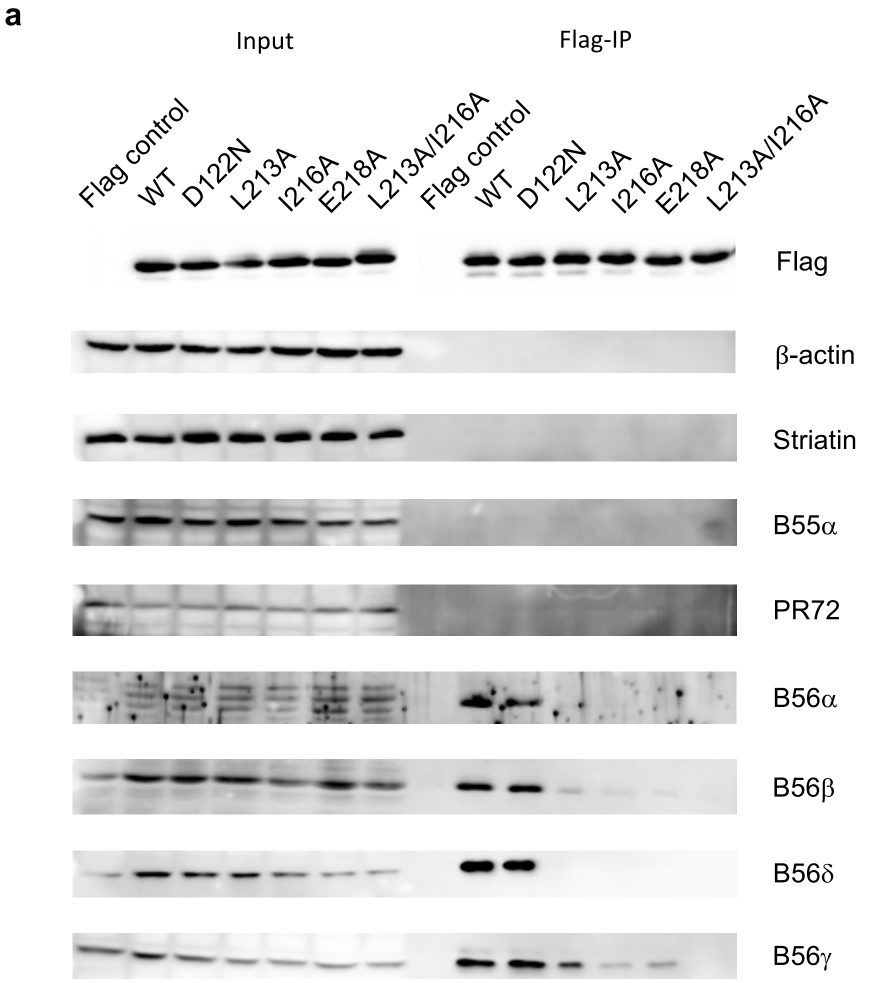


b

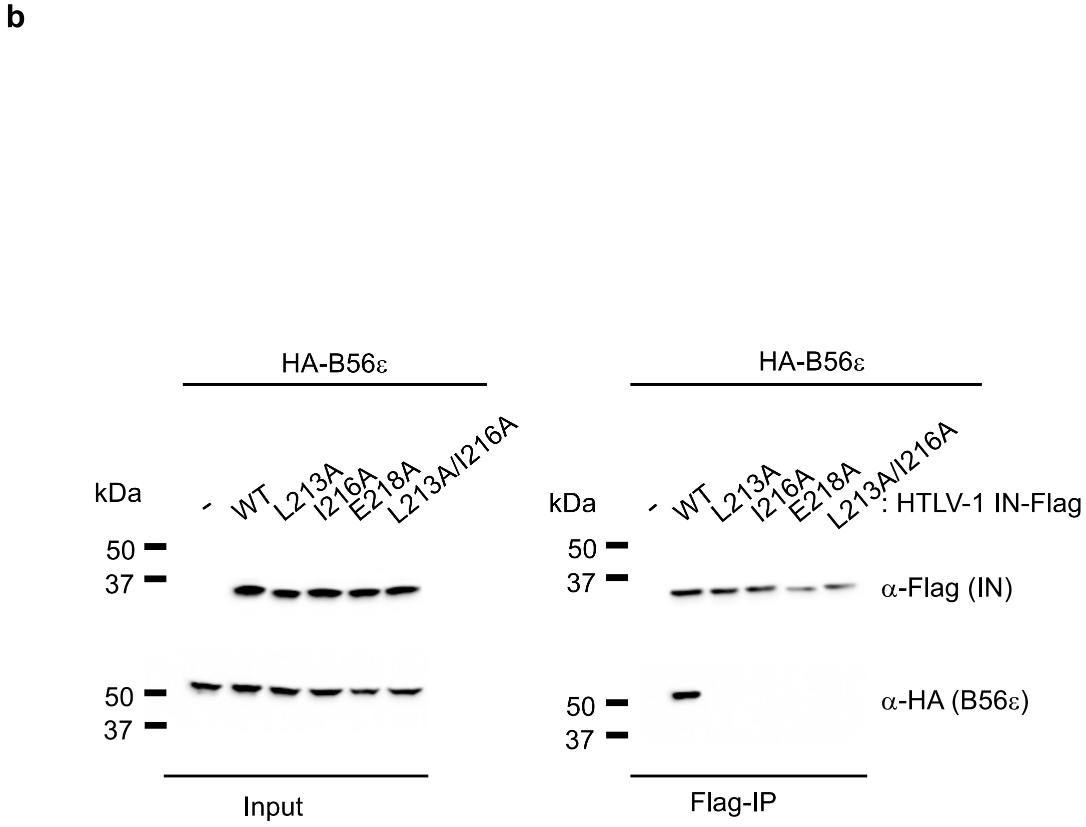

### Supplementary Figure S3_Minnell

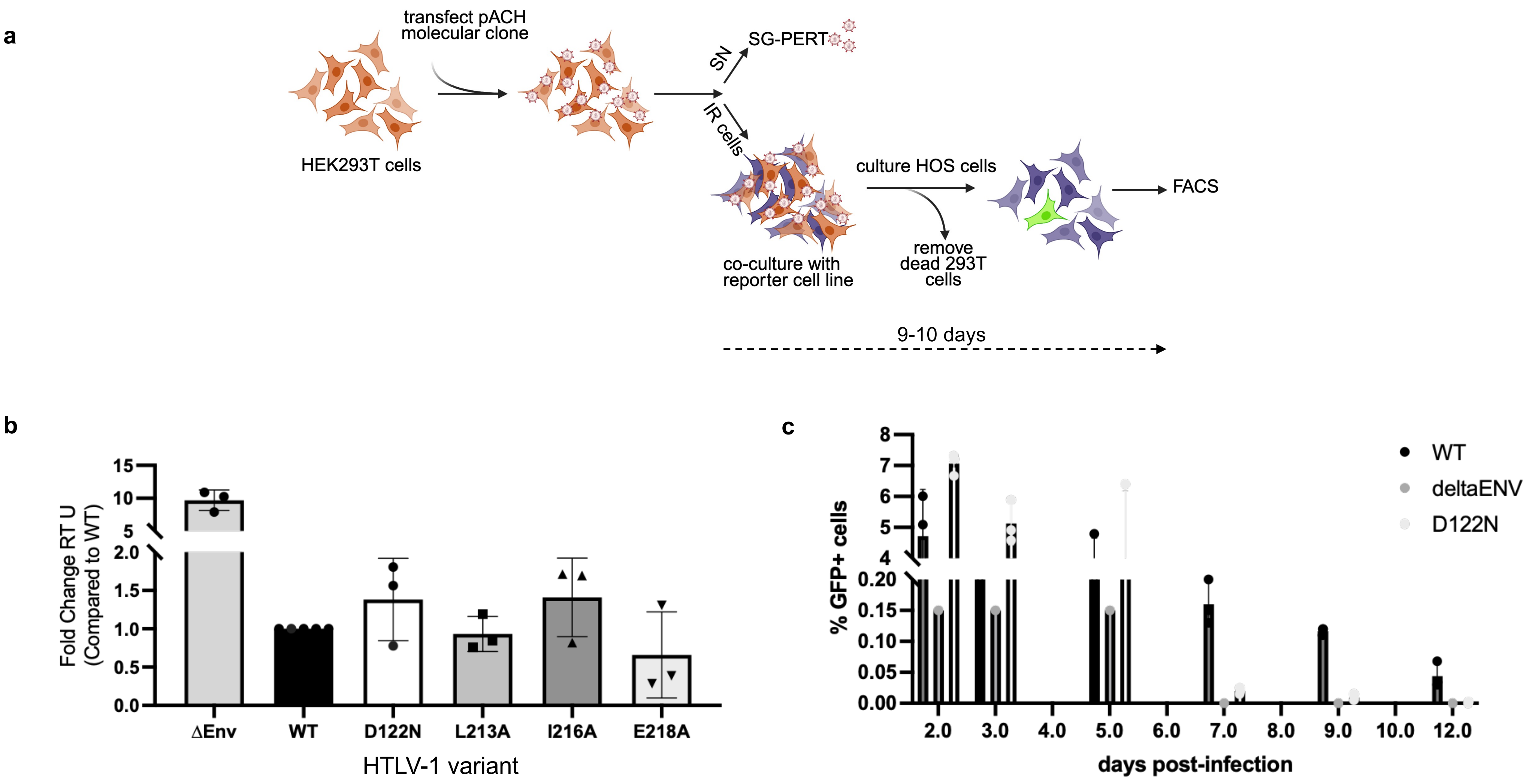

#### Supplementary Figure S4_Minnell

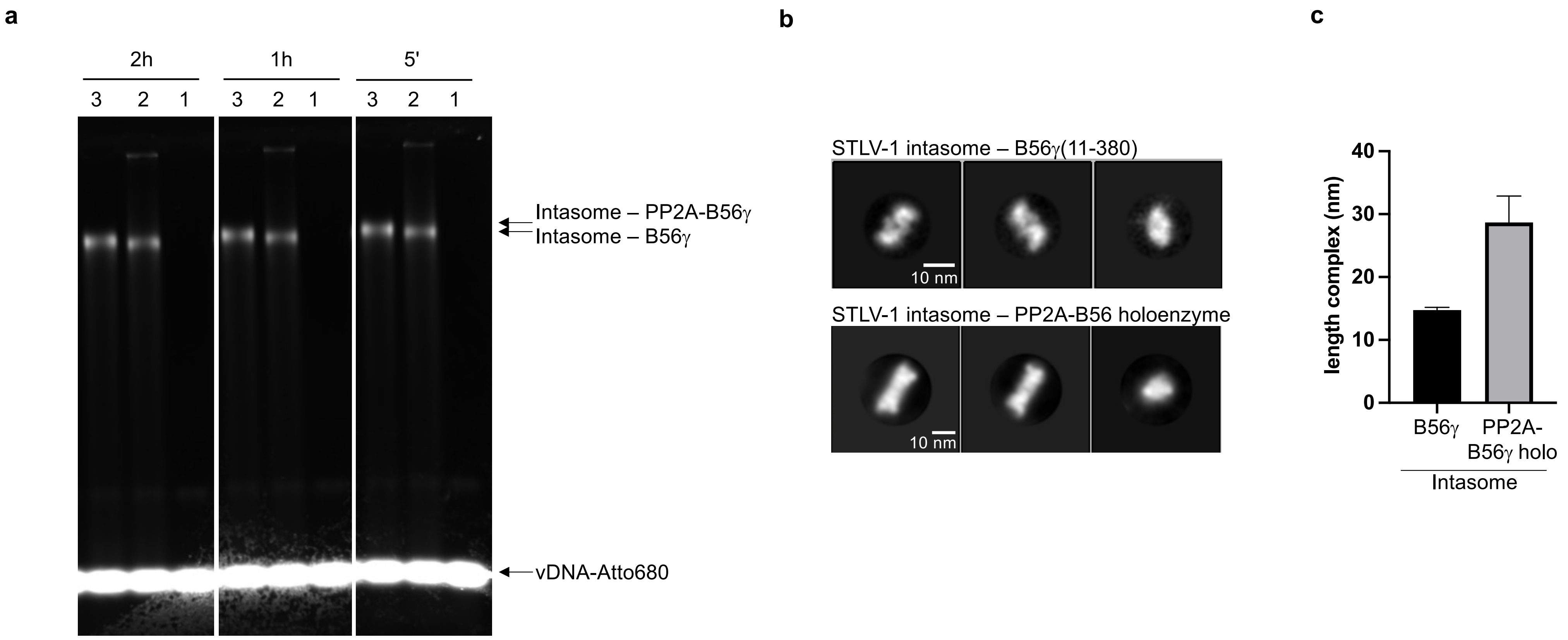

### Supplementary Figure S5_Minnell

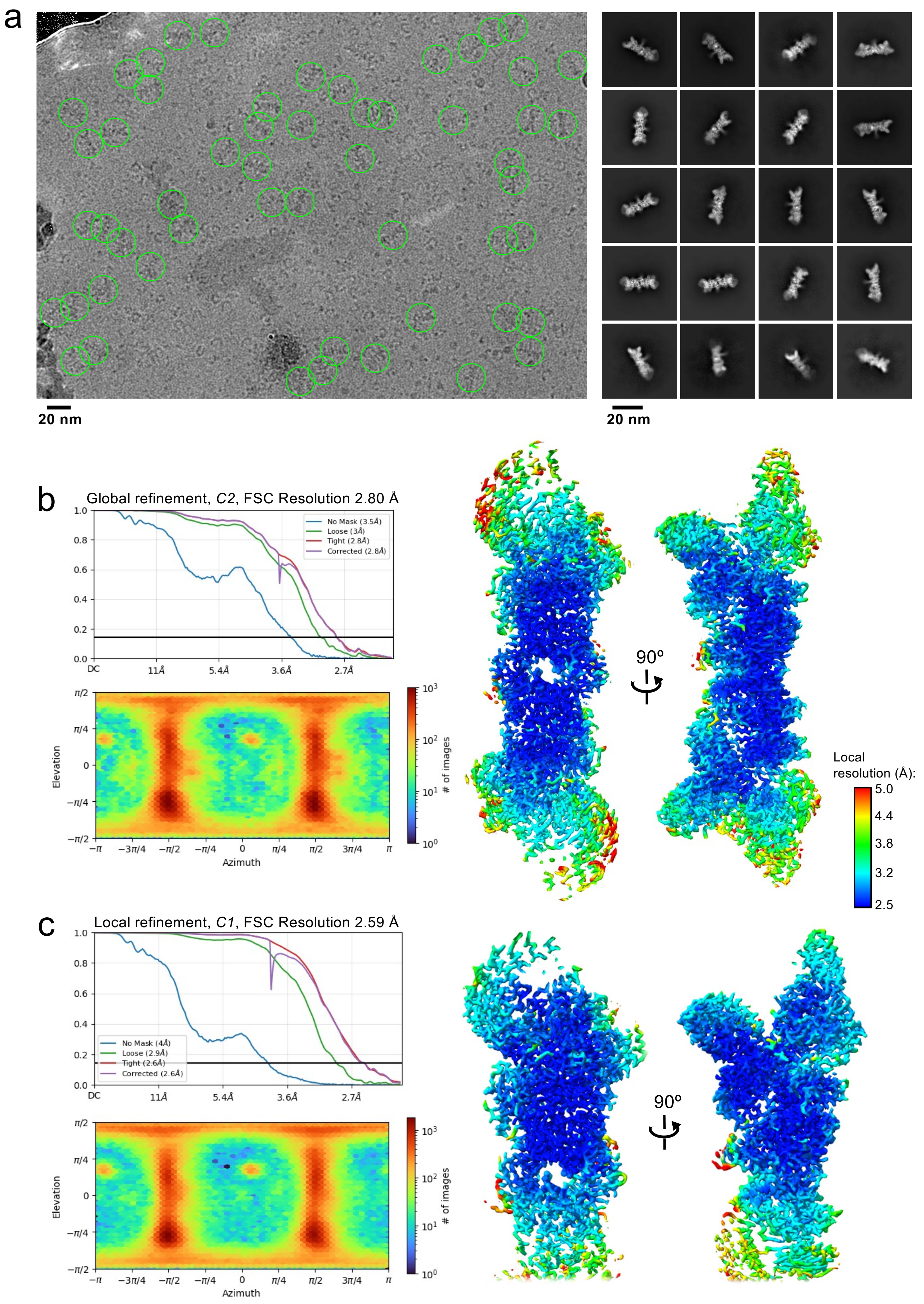

### Supplementary Figure S6_Minnell

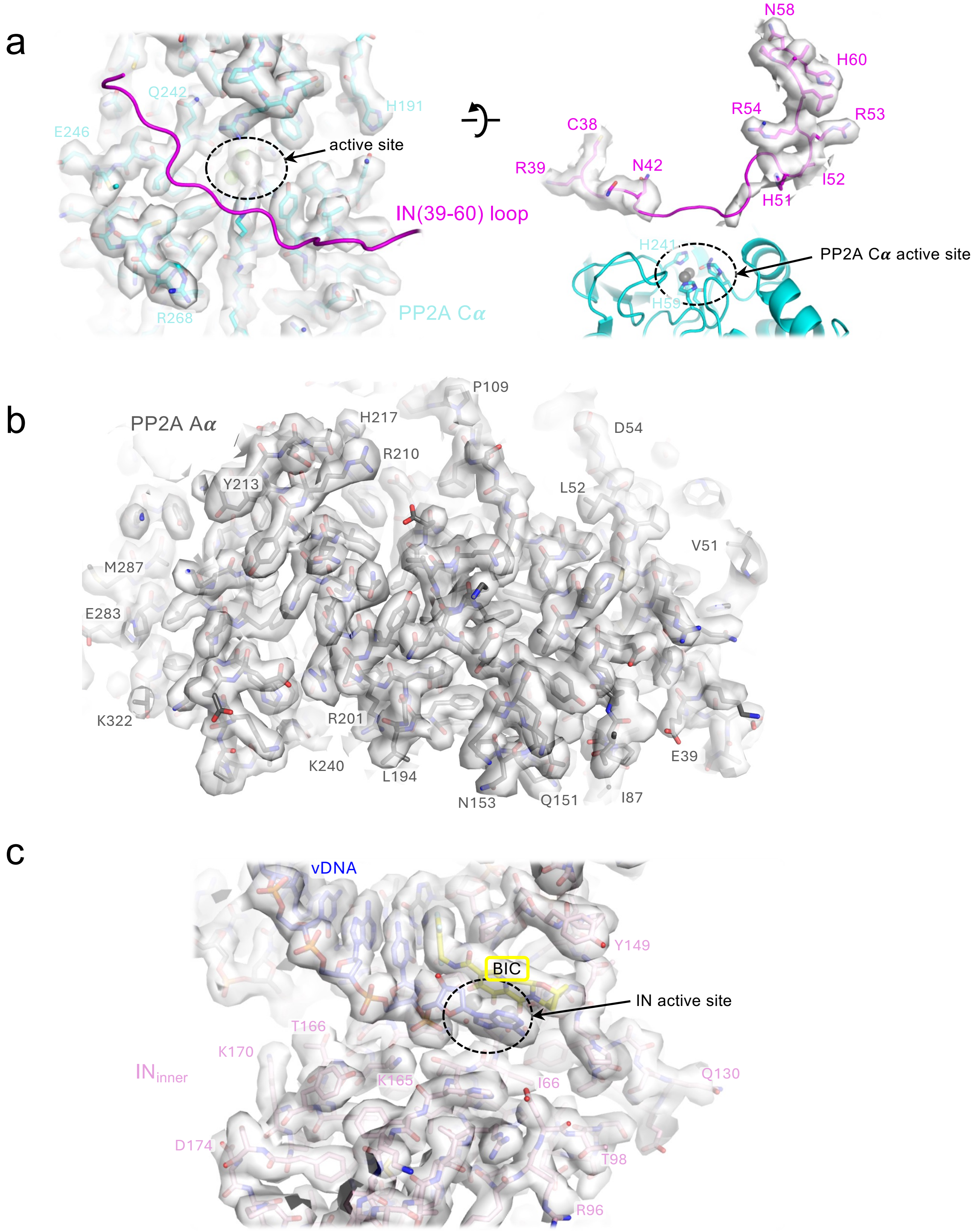

**Supplementary Figure S7_Minnell**

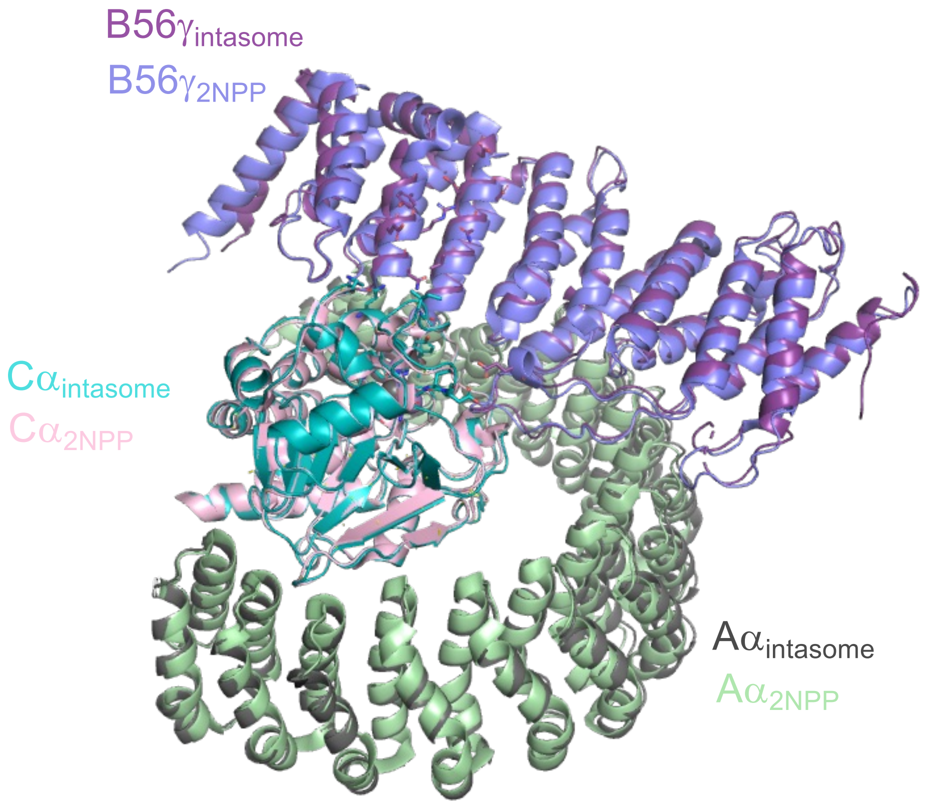

### Supplementary Figure S8_Minnell

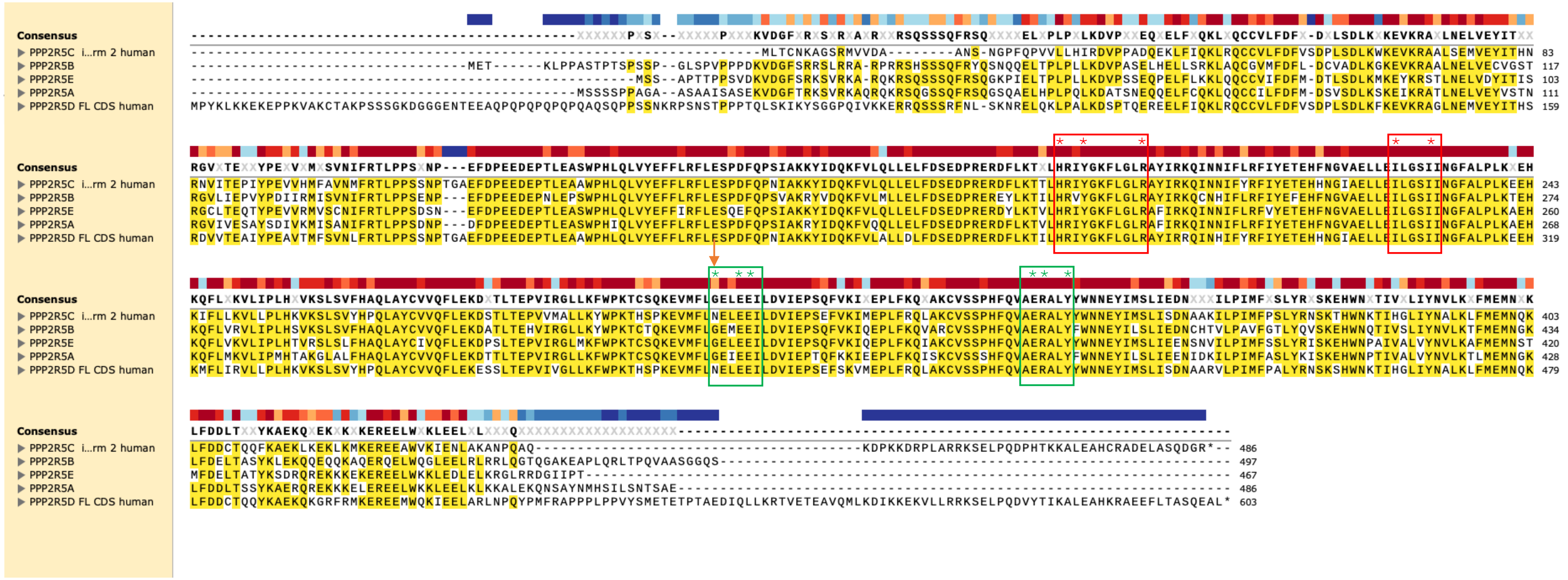
